# Homogeneous selection promotes microdiversity in the glacier-fed stream microbiome

**DOI:** 10.1101/2020.12.03.409391

**Authors:** Stilianos Fodelianakis, Alex D. Washburne, Massimo Bourquin, Paraskevi Pramateftaki, Tyler J. Kohler, Michail Styllas, Matteo Tolosano, Vincent De Staercke, Martina Schön, Susheel Bhanu Busi, Jade Brandani, Paul Wilmes, Hannes Peter, Tom J. Battin

## Abstract

Microdiversity, the organization of microorganisms into groups with closely related but ecologically different sub-types, is widespread and represents an important linchpin between microbial ecology and evolution. However, the drivers of microdiversification remain largely unknown. Here we show that selection promotes microdiversity in the microbiome associated with sediments in glacier-fed streams (GFS). Applying a novel phylogenetic framework, we identify several clades that are under homogeneous selection and that contain genera with higher levels of microdiversity than the rest of the genera. Overall these clades constituted ∼44% and ∼64% of community α-diversity and abundance, and both percentages increased further in GFS that were largely devoid of primary producers. Our findings show that strong homogeneous selection drives the microdiversification of specialized microbial groups putatively underlying their success in the extreme environment of GFS. This microdiversity could be threatened as glaciers shrink, with unknown consequences for microbial diversity and functionality in these ecosystems.

## Introduction

Microdiversity, the organization of microbial sub-taxa with distinct niches within a larger phylogenetic clade^1^, is an intrinsic property of microbial communities that is widely distributed in nature. Whereas initial studies suggested that microdiversity results from genetic drift and does not necessarily involve phenotypic differences^2,3^, such differences are now well documented in major biomes of the planet. In marine environments, microdiversity within marine picocyanobacteria (e.g., *Prochlorococcus, Synechococcus*) and heterotrophic bacteria (e.g., *Vibrio, Pelagibacter)* includes sub-taxa acclimated to light intensity, nutrient availability and hydrostatic pressure^4-9^. In freshwaters, members of the ubiquitous *Limnohabitans* exhibit microdiversity related to temperature and nutrient availability^10^, while microdiversity has also been reported from *Streptomyces* and *Curtobacterium* in soils along climatic^11^ and latitudinal^12^ gradients, respectively.

Microdiversity results in phenotypic (i.e., trait) differentiation, and traits affect the fitness and performance of a taxon under specific environmental conditions^1,13^. Therefore, from a community ecology perspective, it is intuitive to assume that microdiversity relates to the assembly process of selection, i.e., deterministic fitness differences among species^14^. Recent microbial community surveys tend to support this notion, for instance by observing different temporal turnover patterns among marine sub-taxa^15,16^ or by examining the distribution of microbial sub-taxa along environmental gradients^17^ and their participation in different biotic interactions^18^. However, it is yet unclear whether there is a causal relationship between the community assembly process of selection (henceforth referred to as “selection”) and microdiversity (i.e., whether selection promotes microdiversity). Here we posit that such a relationship exists between microdiversity and selection, the latter being expanded to include fitness differences *within* species as well, given that bacterial species boundaries are blurry^19^. More specifically, we hypothesize that this relationship should be more prevalent in ecosystems where community assembly is largely driven by selection. Under the assumption of phylogenetic conservatism (i.e., closely related taxa share similar phenotypes), taxa with high fitness in such ecosystems should form coherent phylogenetic clades. Within these clades, closely related taxa with a specific fitness advantage may occupy diverse niches via fine-tuning or gain of novel phenotypic traits^1^. Eventually, this would foster the generation of microdiversity within clades under selection. Hence, empirical evidence showing a relationship between clades under selection and microdiversity in an ecosystem dominated by selection would suggest that selection fosters microdiversification.

Phylogenetic clades under selection could be identified by expanding the analytical framework that is used to infer community-wise assembly processes^20,21^. At the community level, lower phylogenetic turnover than expected by chance indicates communities under homogenous selection^20,21^. Homogeneous selection indicates that the same selective processes (e.g., abiotic conditions or biotic interactions) are acting among the studied communities, and it appears to dominate community assembly in extreme, energy-limited ecosystems^22-24^. In analogy, the presence of phylogenetically similar sequence variants (SVs) occurring more frequently than expected by chance could be used to identify phylogenetic clades under homogeneous selection.

Glacier-fed streams (GFS) seem well suited to test the potential relationship between microdiversity and selection, because of their extreme environment (e.g., low temperature, oligotrophic conditions, hydraulic stress) and diverse communities of microbes and macroorganisms^25-27^. We expect that because of the extremeness in the GFS environment, homogenous selection results in the prevalence of a few highly specialized phylogenetic clades that are characterized by enhanced microdiversity. To that end we studied sediment biofilms at twenty GFS across a 340-km long transect in the Southern Alps in New Zealand (Fig. S1). Our sampling design allowed us to capture patterns of community turnover over a large spatial scale as well as within streams (two reaches per stream and three biological triplicates per reach) where dispersal should be more important and could potentially attenuate selection with mass effects^21,28^ via the water flow. Subsequently, we quantified the community assembly processes, developed a novel framework to identify phylogenetic clades under selection and examined if they are characterized by a high degree of microdiversity.

## Results

### Homogeneous selection is the dominant assembly process at the community level

Using a community-level framework^20,21^, we first examined the processes that govern the assembly of sediment biofilm communities among and within the GFS (*Methods*). We found that homogeneous selection (reflected as βNTI values < -2) was the dominant assembly process for 89.2% of the community pairs among GFS (Fig. 1). Moreover, homogeneous selection dominated (in 99.3% of the community pairs) the assembly within GFS, indicating that it was not attenuated by downstream dispersal via water flow. Dispersal limitation drove assembly for 9.5% of community pairs among GFS and its probability of occurrence increased with increasing geographic distance between the compared communities (logistic regression, z=11.97, p < 0.001). Finally, variable selection and homogenizing dispersal drove assembly for 0.6% and 0.25% of community pairs among GFS, respectively, while no single dominant process was found in 0.45% of community pairs.

**Figure 1.**
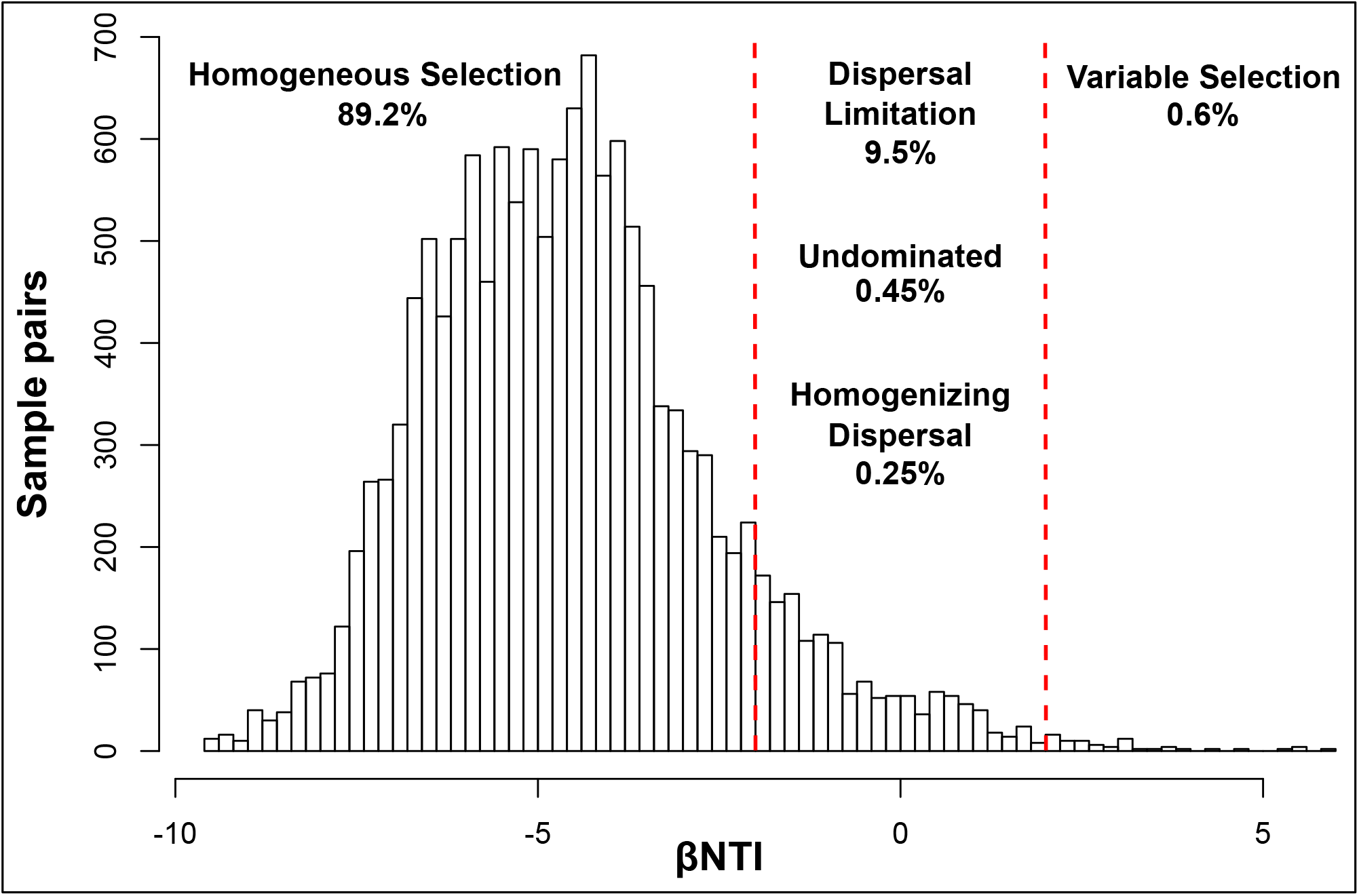
Homogeneous selection is the dominant assembly process at the community level. The histogram shows the distribution of βNTI values for sample comparisons across GFS and the proportion of sample pairs under each community assembly processes. Vertical dashed red lines are drawn at βNTI values of -2 and +2, which are the cutoff values for lower and higher than expected phylogenetic community turnover, respectively, the former indicating homogeneous selection and the latter indicating variable selection. The assembly processes governing the sample pairs in between are estimated from compositional turnover patterns based on the RC_Bray_ index.

### Phylogenetic clades under homogeneous selection are diverse, abundant and widespread

Next, having confirmed the dominant role of selection in driving assembly at the community level, we developed and applied a method that leverages null phylogenetic modeling to identify phylogenetic clades that are under homogeneous selection. In analogy to the community-level framework, we defined these clades as phylogenetically coherent groups that contain SVs with phylogenetically closer relatives across communities than expected by chance. To identify such SVs, we used a z-score that describes how phylogenetically distant an SV in one community is to its closest relative in another community compared to what is expected by chance. Specifically, the z-score counts this phylogenetic distance in standard deviations with respect to a null distribution of phylogenetic distances so that negative z-scores represent shorter phylogenetic distances than expected by chance and *vice versa*. We then calculated the total z-scores for each SV across the dataset, i.e., the sum of the z-scores of that SV across all community pairs, excluding sample pairs from replicates of the same reach. For a given SV, highly negative total z-scores indicate that it is replaced by closely related SVs in many community pairs. In the presence of a phylogenetic signal at short phylogenetic distances, which we verified for our dataset (*Methods*, Fig. S2), this indicates that the given SV has increased fitness in the specific ecosystem, because it and its functionally similar close relatives are widespread. Consequently, we used phylofactorization^29,30^ to identify phylogenetically coherent groups of SVs that have significantly different total z-scores compared to outgroups.

We identified eight phylogenetic clades with significantly lower total z-scores compared to outgroups (contrast tests, 3.3E-16 < p < 6.8E-255), comprising of 5 to 1418 SVs each (Fig. 2, Table S1). The consensus taxonomy of the largest identified clade (1418 SVs) affiliated to *Betaproteobacteria* (Fig. 2). This clade also contained three sub-clades with distinctly low scores and with consensus taxonomies affiliated to the family *Comamonadaceae* (575 SVs), to the uncultured order Ellin6067 (54 SVs) and to the genus *Methylotenera* (48 SVs). The second largest clade (602 SVs) had a consensus taxonomy affiliated to *Alphaproteobacteria* and it contained a low-score sub-clade (5 SVs) affiliated to the genus *Novosphingobium*. The third largest clade (338 SVs) was affiliated to the candidate class *Saprospirae* within *Bacteroidetes* while the smallest clade (18 SVs) was taxonomically affiliated to the genus *Nitrospira*. Importantly, we did not identify any phylogenetic clade with significantly higher total z-scores than expected by chance; this reflects the low contribution of heterogeneous selection in governing assembly at the community level (i.e., low percentage of community pairs with higher than expected phylogenetic turnover; Fig. 1).

**Figure 2.**
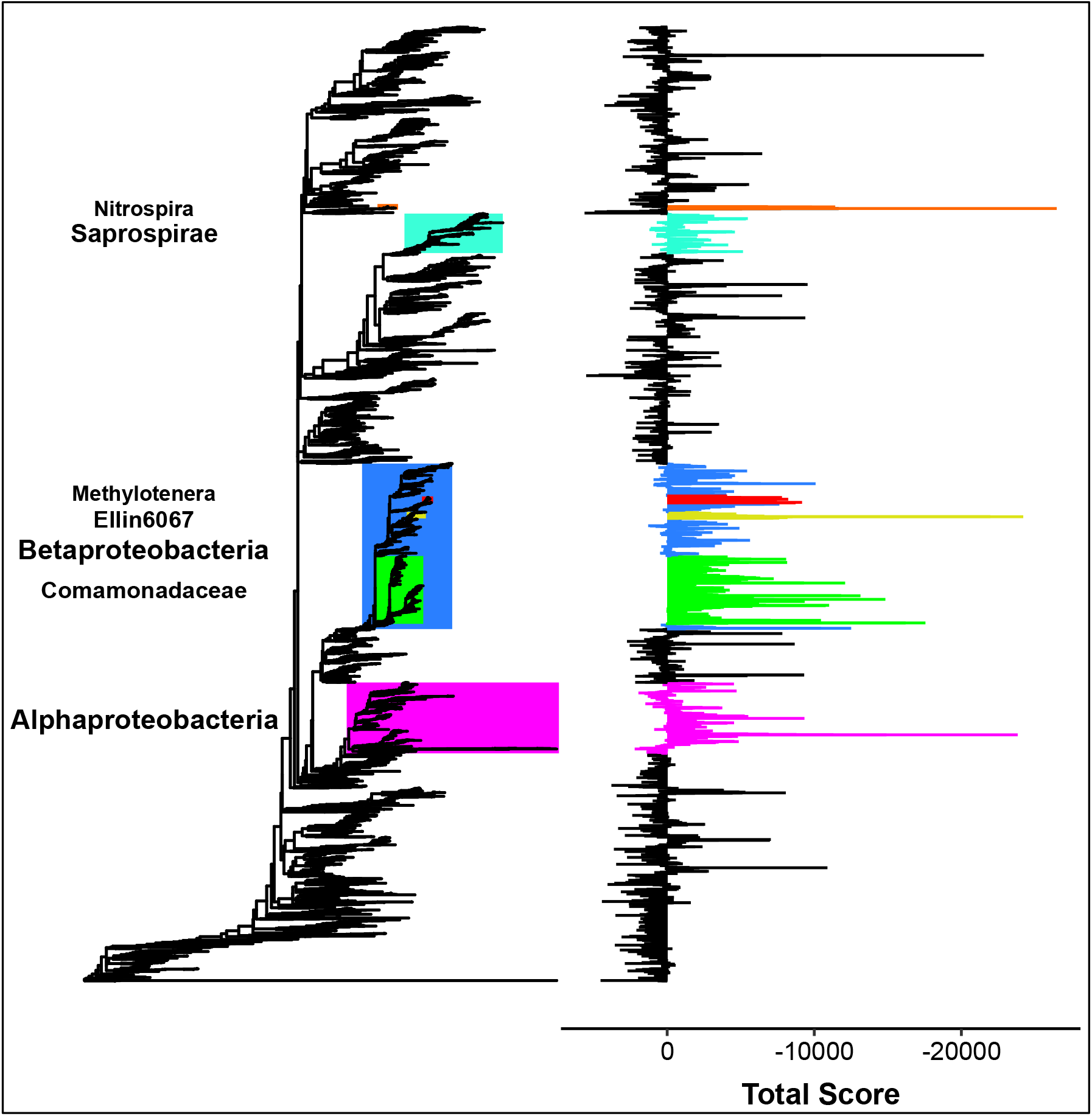
The identified phylogenetic clades have lower within-clade total z-scores compared to outgroups. Identified clades with more than 15 SVs are color-coded on the phylogenetic tree, and the consensus taxonomy is given for each clade on the left with font size proportional to taxonomic depth. Clades nested within Betaproteobacteria are colored individually. The total score of each SV (i.e., the sums of the z-scores across community pairs) is shown to the right as bars with colors matching the clades’ colors.

We found that these clades contained a significant part of the total bacterial diversity and abundance at all GFS, with on average 43.7% (25.5% to 61.6%) of the total SVs and 64% (37.6 to 83.3%) of the total sequences per sample. Additionally, there was a notable overlap between the identified clades and the core microbiome defined as the bacterial genera present at all reaches (Table S2, Supplementary Results). More specifically, nine of the twelve core genera resided within the identified phylogenetic clades (Fig. 3); these genera included the majority of the SVs (59.5%) and of the sequences (87.7%) present in the core genera. Furthermore, both the abundance and the α-diversity of the identified clades increased disproportionately compared to the rest of the microbiome as sediment chlorophyll *a* decreased (linear models, n=119, adjusted R^2^ = 0.298 and 0.302, respectively, and p< 0.001 for both models) (Fig. 4A-B). Since sediments with lower chlorophyll *a* also contained fewer total bacterial cells (Pearson correlation, r=0.85, p<0.001) (Fig. 4C), the above correlations held true with decreasing cell numbers as well (linear models, n=119, adjusted R^2^ = 0.282 and 0.252, respectively, and p< 0.001 for both models) (Fig. S3). Collectively, these results indicate that the identified clades are ecologically successful in the extreme GFS environment, underscoring their apparent fitness and corroborating that they are under homogeneous selection.

**Figure 3.**
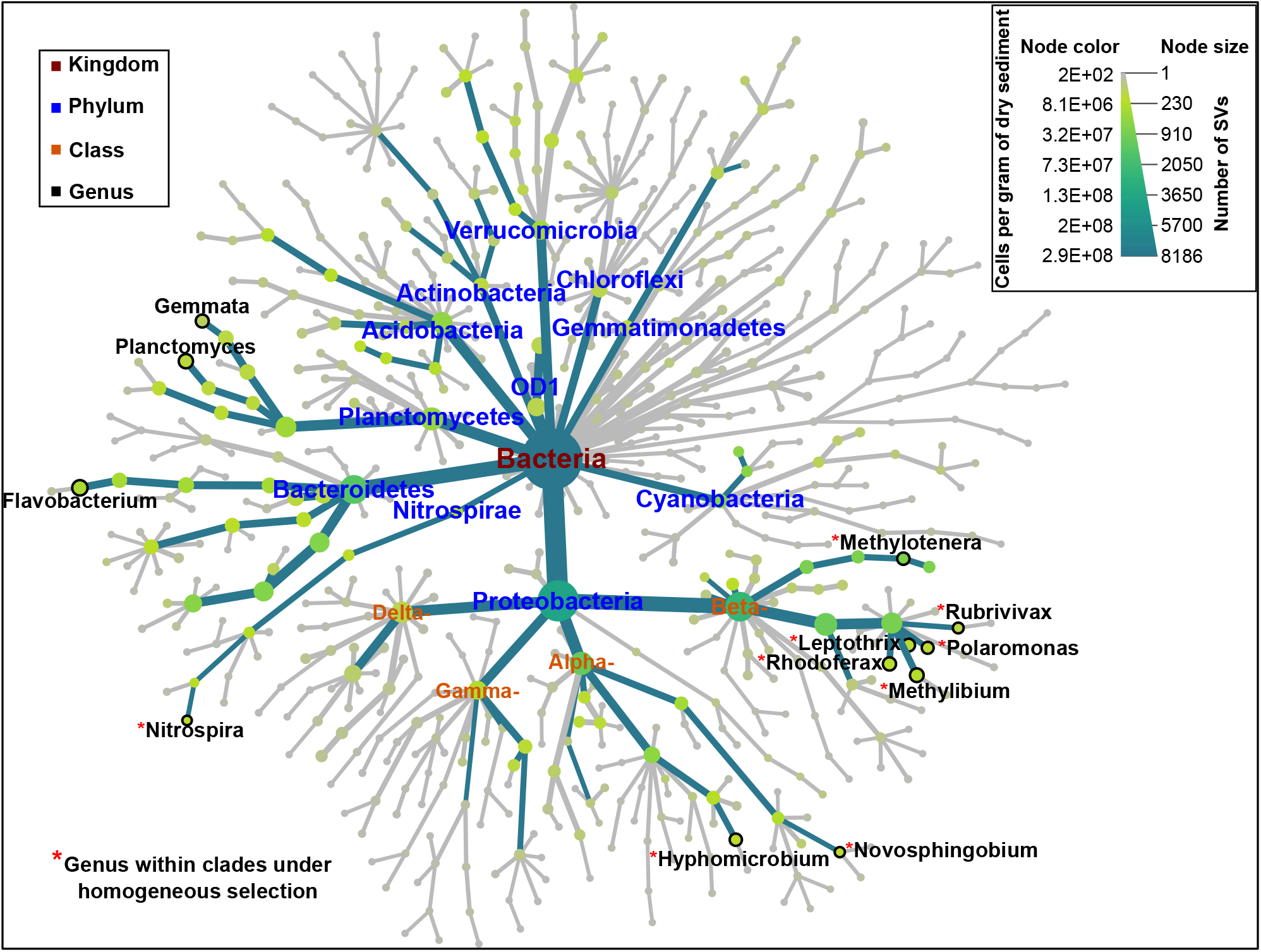
The core microbiome at the genus level and the phylogenetic clades under homogeneous selection overlap highly. The overall core microbiome, i.e., taxonomic units found across all the sampled GFS reaches, is represented as a hierarchy tree (dark green edges) within the overall taxonomic tree (dark green and grey edges). The node color and size are proportional to the node’s abundance (cells per gram of dry sediment) and diversity (number of SVs), respectively, as per the legend on the upper right. Only core genera, phyla, and classes within the Proteobacteria phylum are labeled to improve visualization with colors according to the legend on the upper left. Red asterisks indicate genera that reside in phylogenetic clades under homogeneous selection.

**Figure 4.**
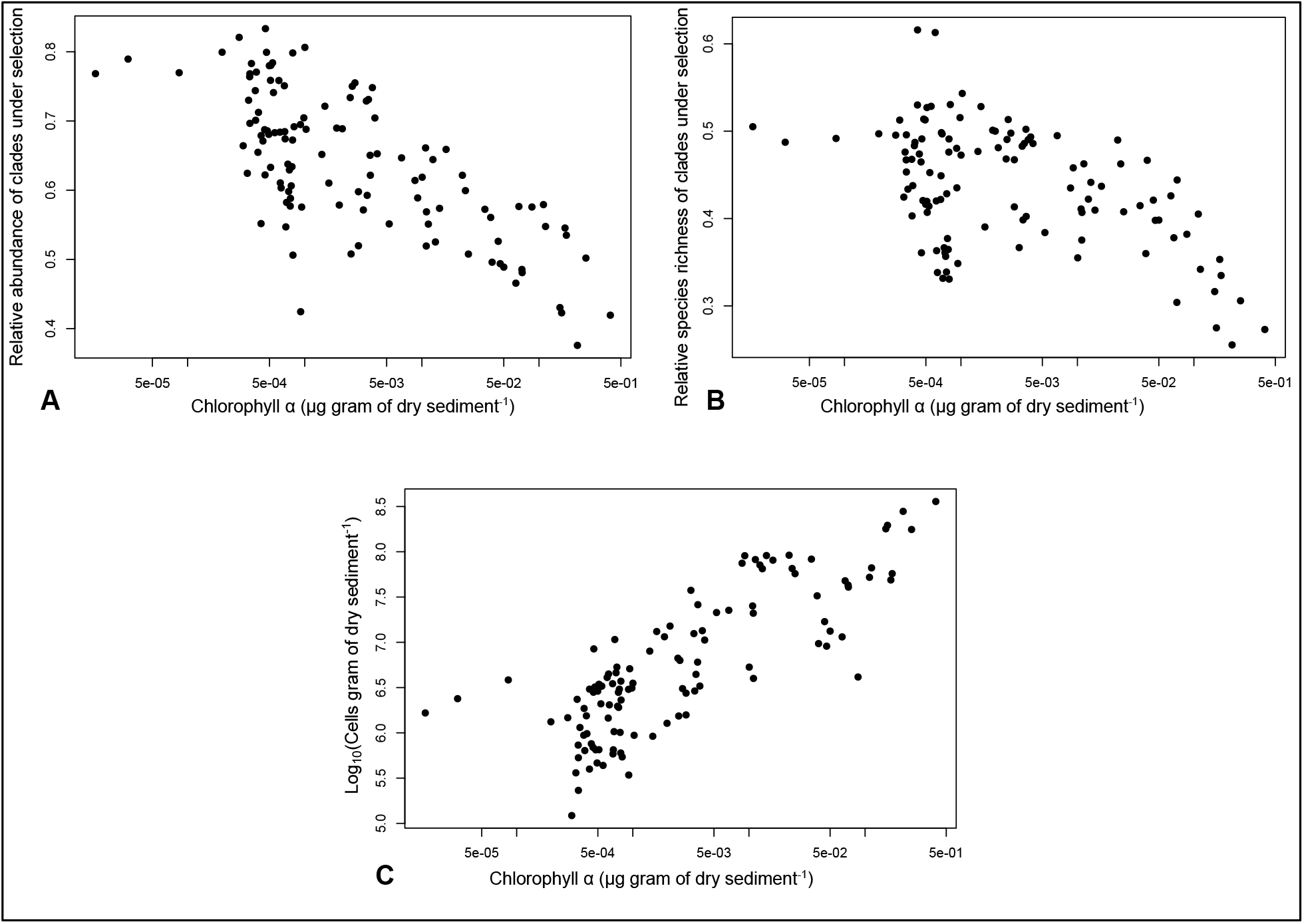
The phylogenetic clades under homogeneous selection thrive in sediments with low chlorophyll *a* that also have low total bacterial cell abundance. **A**. The relative abundance of the clades under homogeneous selection as a function of the sediment chlorophyll *a*. **B**. The relative species richness of the clades under homogeneous selection as a function of the sediment chlorophyll *a*. **C**. The total bacterial cell abundance as a function of the sediment chlorophyll *a*. For all panels, n=119.

### Eleven genera within phylogenetic clades under homogeneous selection are hotspots of microdiversity

Having identified the phylogenetic clades under homogeneous selection, we next explored whether they exhibit disproportionally high levels of microdiversity supported by the occurrence of numerous, closely related and ecologically distinct subtaxa. To that end, we compared similar taxonomic units, assessing the levels of microdiversity in genera within and outside these clades. We restricted our analyses to genera with at least 2 SVs that are formally assigned in the scientific nomenclature, resulting in the inclusion of 110 genera containing 2003 SVs in total. This represented approximately half (47.2% on average, 26.5-79% per sample) of the total sequences. Phylogenetic clades under homogeneous selection included 41 genera, whereas 69 genera resided outside these clades (Table S3). Within-genera, microdiversity should emerge as high numbers of closely related SVs. Moreover, if closely related SVs within genera indeed occupy distinct niches, this should result in a wide genus spatial distribution. We compared the following three attributes of all genera: the number of SVs per genus, the mean nucleotidic similarity and the spatial distribution (i.e., the B parameter *sensu* Levins^31^) as a proxy for niche breadth given that there was no single environmental gradient driving β-diversity (*Methods*) (Supplementary Results, Fig. S4).

Our results revealed eleven genera, namely *Methylotenera, Rhodoferax, Leptothrix, Polaromonas, Methylibium, Rubrivivax, Thiobacillus, Novosphingobium, Hyphomicrobium, Rhodobacter* and *Nitrospira*, with disproportionately higher levels of microdiversity compared to other genera (Fig. 5). Strikingly, all these genera resided within clades under homogeneous selection and represented a large part of the SVs (33%) and of the cells (average: 55.5%; range: 35.1-78%) therein. Furthermore, these genera had high numbers of SVs per genus (average: 71.9; range: 20-130), as well as high mean pairwise nucleotidic similarity (average: 96.1%; range: 95.1-96.7%) and large mean niche breadths (average: 5.96; range: 4.62-8.12) (Fig. 5). Specifically, they had approximately 10 to 50-fold higher numbers of SVs per genus compared to other genera, as well as higher mean pairwise nucleotidic similarity (t-tests, p<<0.001 for all comparisons) and niche breadths (t-tests, 6.1E-05 < p < 0.04) compared to the species-rich genera of *Flavobacterium, Planctomyces, Gemmata* and *Bdellovibrio* that do not reside within phylogenetic clades under homogeneous selection (Fig. 5). Overall, our results show that the eleven identified genera were the only genera across the dataset that simultaneously had high numbers of SVs (per genus), high mean pairwise nucleotidic similarity and wide spatial distribution. This is evidence that microdiversity is disproportionally more present within phylogenetic clades under homogeneous selection.

**Figure 5.**
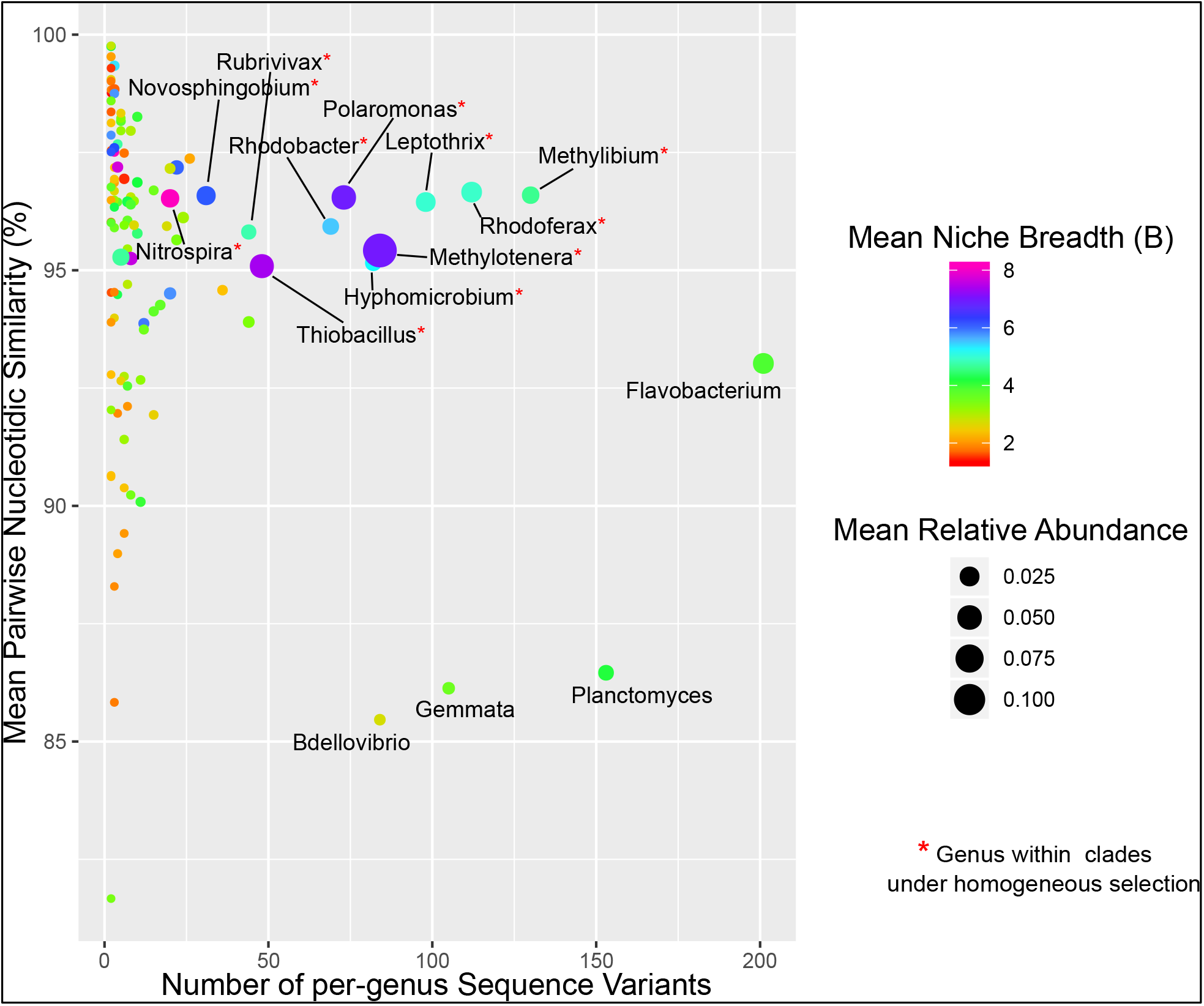
Microdiversity is disproportionately more present in genera residing within phylogenetic clades under homogeneous selection. The plot shows the per-genus degree of microdiversification, assessed via the number of per-genus Sequence Variants (SVs), the mean pairwise nucleotidic similarity and the mean niche breadth (B – color coding). Dot sizes are proportional to the mean relative abundance. Labels are shown only for genera with more than 50 SVs, or for the eleven identified genera within phylogenetic clades under selection, the latter labeled with an additional red asterisk. Note the high number of per-genus SVs of *Flavobacterium, Planctomyces, Gemmata* and *Bdellovibrio*, which do not reside within clades under homogeneous selection.

### SVs with low phylogenetic turnover foster microdiversity within specific genera

Finally, we asked whether the eleven genera with elevated microdiversity are also characterized by disproportionately low phylogenetic turnover (indicative of strong homogeneous selection). In other words, if selection promotes microdiversity, the eleven genera with the highest levels of microdiversity should also show the strongest signs of homogeneous selection. More specifically, the eleven genera with the highest levels of microdiversity should contain clusters of SVs with low phylogenetic turnover that are driving the increase of microdiversity within the genus.

To test that, we first detected SVs with constantly low phylogenetic turnover (LPT-SVs) and their phylogenetically closer-than-expected relatives (CR), to then examine if they are primarily present within the eleven genera of interest. Subsequently, we examined if these clusters are also characterized by high sequence similarity and high niche differentiation. We defined LPT-SVs as SVs that have lower z-scores than expected by chance in the majority of the applicable community comparisons (SVs having a median z-score < -2). LPT-SVs represent SVs with a CR across most GFS and do not necessarily have high occupancies. For example, an LPT-SV of the genus *Methylibium* in our dataset is only present in four reaches across all GFS, but has three CRs (with which it is 99.7% similar at the sequenced part of the 16S rRNA gene on average) in 204 out of the 344 applicable community pairs that span 33 out of the 40 reaches across all GFS. In other words, these four SVs (the LPT-SV and its CRs) are highly similar genetically and have a broad spatial distribution as a cluster but not individually. Taking spatial distribution as a proxy for niche breadth and having already excluded the presence of dispersal limitation, this pattern points towards the coverage of distinct niches by these sub-taxa. Therefore, a strong presence of LPT-SVs in the eleven genera would indicate that these genera broaden their spatial distribution in GFS by microdiversification.

Indeed, we found that the clusters of the LPT-SVs and their CRs were disproportionally present in the eleven genera and they had high genetic similarity and wide spatial distributions. We identified a total of 172 LPT-SVs and found that their majority (143 SVs) resided within phylogenetic clades under homogeneous selection. Furthermore, these LPT-SVs had very low phylogenetic turnover, accounting for 80.5% of the total z-score among LPT-SVs (Table S4). Importantly, 104 (60.5%) of the LPT-SVs were taxonomically assigned to the eleven genera of interest with nine out of these genera containing at least one LPT-SV (Table 1). The LPT-SVs and their CRs had a mean nucleotidic similarity of 97.2-99.8%, which was higher than that of the respective genus. This indicates that they represent clusters of sub-taxa. These clusters had broad spatial distributions, occupying 32 to 40 of the 40 total reaches across all GFS. In most of the cases, except in *Methylibium* and in *Rhodobacter* that contained only 3 and 1 LPT-SV, respectively, the reach occupancy of LPT-SVs and CRs was higher than that of the rest of the SVs within the same genus. This indicates that the former contributes in widening the spatial distribution of the respective genus. Moreover, LPT-SVs and CRs were highly abundant within the respective genus (except in *Methylibium* and *Rhodobacter*), comprising 47.2-99% of the total cells affiliated with this genus. These results confirm that clusters of SVs with low phylogenetic turnover are driving the increase of microdiversity in the eleven genera of interest, and suggest that homogeneous selection promotes microdiversity in the examined GFS sediments.

**Table 1.**
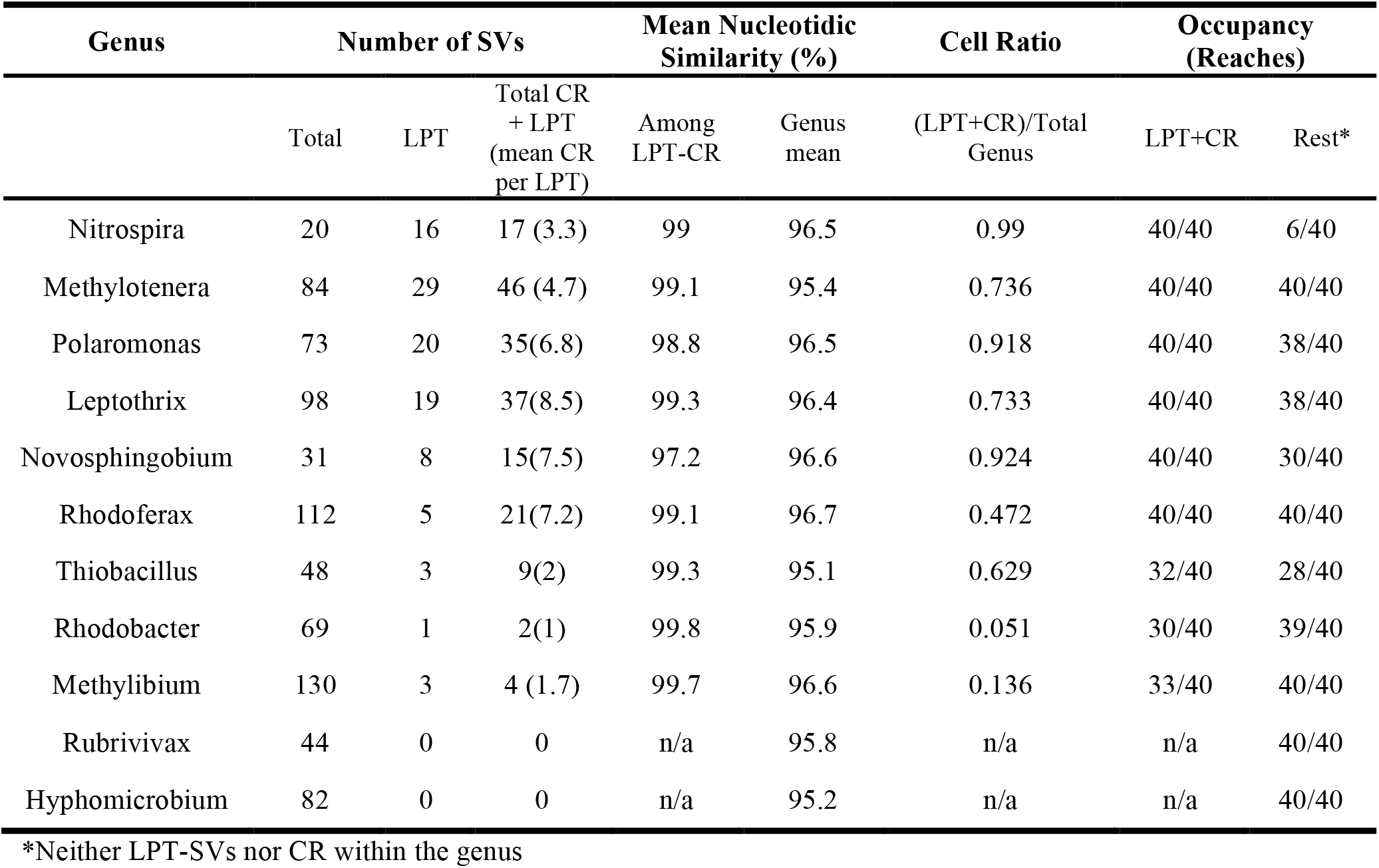
The presence and properties of low phylogenetic turnover Sequence Variants (LPT-SVs) and their phylogenetically closer than expected relatives (CR) in the eleven genera that reside within clades under homogeneous selection and that have higher levels of microdiversity compared to other genera.

## Discussion

Employing a novel analytical framework for phylogenetic turnover analysis, we showed that selection promotes microdiversity in the microbiome associated with the bed sediments of GFS. Low phylogenetic turnover, attributed to homogeneous selection in community ecology^20,21^, dominated the assembly of microbial communities as is typical for energy-limited environments^22-24^. Our analytical framework further allowed us to dissect the contribution of individual phylogenetic clades to this low phylogenetic turnover. In analogy, we call these clades as being under strong homogeneous selection, and both their high occupancies and abundances in GFS corroborate this notion. Focusing on the eleven genera within these clades with high microdiversity, we found that microdiversity was fostered by specific fine-scale clusters of SVs with low phylogenetic turnover. These findings shed new light on the drivers of the fine-scale phylogenetic architecture and its consequences for the success of microbial life in the extreme GFS environment.

Our results further suggest that the phylogenetic clades under homogeneous selection can successfully occupy a niche in the GFS that is largely devoid of algal primary producers. This is indicated by their stronger presence in sediments with little chlorophyll *a*. Concomitantly, lower cell abundance in these sediments further evokes that the rest of the microbiome is energy-limited in these sediments. We interpret these patterns as evidence for an ecological niche governed by chemolithotrophic rather than heterotrophic energy pathways, as is typical in extreme environments, including the cryosphere and deep biosphere^32-35^.

This notion is indeed supported by the physiologies broadly studied and established for the genera residing within the clades under homogeneous selection. For instance, the globally-spread^36^ psychrophilic genus *Polaromonas* is facultatively chemolithotrophic and metabolically versatile^37^, and was even reported to be microdiverse^38^. Furthermore, the obligate methylotrophs *Methylibium, Methylotenera* and *Hyphomicrobium* have been found in deglaciated alpine soils^39^ and glaciers^40^, and can utilize a diverse array of C_1_ compounds^41-43^ that can occur as intermediates in methane oxidation typical for the sub-glacial environment ^44,45^. The anoxygenic phototrophs and nitrogen fixing genera *Rhodobacter, Rubrivivax* and *Rhodoferax* include psychrotolerant isolates^46,47^ and have been found in ice cores^48^, deglaciated soils^49^ and glaciers^40,50^. Furthermore, members of the *Nitrospira* genus are ubiquitous nitrite oxidizers, and species able to perform complete ammonium oxidation have recently been reported in a high-altitudinal and cold-water river^51^. The sulfur-oxidizing, facultative anaerobe and chemolithotrophic *Thiobacillus* has a sequenced genome from a subglacial isolate revealing cold adaptations^52^ and is frequently found in cold-related environments^53,54^. The only “classical” heterotroph among the identified genera is the iron oxidizing *Leptothrix*^55^, which has been recently reported from a metagenome from Antarctica^56^.

The environment of GFS is predicted to change dramatically as glaciers shrink owing to climate change^25,57^. A recent synthesis has suggested that specialist species that are well adapted to the glacial conditions in GFS are highly threatened by glacier retreat^25^. At the same time, as turbidity decreases in GFS because of reduced discharge and sediment loads, the environment will become more advantageous for primary production^57^. Therefore, the ecological niche with its microdiverse clades that we have identified will most likely vanish with ongoing glacier shrinkage, and with this, a hidden biodiversity that has adapted to the GFS environment and that could even contain unexploited potential for biotechnology^58^.

Apart from highlighting the importance of microdiversity, our study also contributes in understanding the mechanisms controlling community assembly under the umbrella of deterministic and stochastic processes, which is a debated topic in microbial ecology^59,60^. Analytical frameworks detecting and quantifying assembly processes at the community level^20,21,61,62^ have provided useful insights in a great variety of ecosystems^60^. Recently, the focus has expanded to the identification of specific components of the microbiome that underlie community-level assembly processes. For instance, the recent iCAMP^63^ forms phylogenetic bins of taxa, examines their phylogenetic and taxonomic turnover, and assigns the underlying processes governing their turnover. Our analytical framework for identifying phylogenetic clades under selection is conceptually similar to iCAMP and can be used in parallel with it. Like iCAMP, our framework detects clades with distinctly different phylogenetic turnover than that expected by chance. The detected phylogenetic clades do not necessarily need to have low phylogenetic turnover like in the present study; clades with high phylogenetic turnover indicative of heterogeneous selection (i.e., disproportionally present in different sample groups) can be detected as well. Such patterns would indicate clades under selection in specific spatial or temporal subsets depending on the study, and the assembly of the respective community pairs should be governed by heterogeneous selection. Unlike iCAMP, however, our method avoids phylogenetic binning and uses nearest-taxon phylogenetic distances. Both of these methodological attributes can be valuable when examining patterns near the tips of the phylogeny, such as microdiversity, which might not emerge with the use of other metrics^20^. Therefore, our framework, rooted in community^14^ and metacommunity^64^ ecology, offers a novel data-driven avenue that allows the exploration and quantification of the microdiversity architecture of microbiomes without having to rely on isolates as often required previously^9,11,12,65^. Nevertheless, the short amplicon lengths used in most studies might not be adequate to properly resolve the topology at the tips of the phylogeny. While this should have no effect on the identification of phylogenetic clades with distinct z-scores compared to outgroups, it might affect the identification of specific SVs and thus full-length 16S rRNA gene amplicons^66-68^ might be used to construct phylogenetic trees with highly supported topologies near the tips.

Contrary to the early expectations of an ecologically neutral origin of microdiversity arising from genetic drift^3^, the link between selection and microdiversity that we show here suggests that optimization of niche occupancy probably underlies the observed microdiversification in GFS. This notion is supported by the presence of the same phylogenetic groups that have different and microdiverse SVs in different GFS, probably corresponding to the presence of different microniches therein. We conjecture that this could be a phenomenon common to the microbiome of other extreme environments with reduced competitive exclusion that might have been hitherto unrecognized because of the lack of adequate analytical frameworks. The relaxation of the environmental extremeness owing to climate change may change the balance among the selective processes in GFS and that could erode the microdiversity of the GFS microbiome with yet unknown consequences for the overall biodiversity and ecosystem functioning therein.

## Methods

### Sampling

We sampled 20 glacier-fed streams at the Southern Alps in New Zealand along a 340-km North-East – South-West transect (Fig. S1). The selected glaciers clustered in five major head valley systems (Arthur’s Pass, Westland, Mount Cook, Mount Aspiring and Milford Sound. Glacier surface areas ranged between 0.5 to 35 km^2^, so that a wide range of runoff conditions was encountered during sampling. We sampled stream bottom sediments from two reaches within each of the 20 glacier-fed streams. The upper reaches (hereafter referred to as UP) were located the closest possible to the glaciers’ snouts. The lower reaches (hereafter referred to as DN) were located 100 to 2500 m downstream, representing a gradient of decreasing connectivity with the UP reaches in the same stream via the water flow. For operational purposes, we assigned numbers to the sampled streams from 1 to 21, skipping number 4 (Fig. S1).

Within each reach, we sampled sediments from three different patches in order to assess the within-reach variability. The patches were distanced 2-5 m apart. We sieved the wet sampled sediments through two overlapping 315 and 250 µm fire-sterilized sieves (Retsch, Woven Wire Mesh Sieve - ø 200 mm / 203 mm). We placed 30 grams of sediment in 10-ml cryovials (VWR) and we flash froze them in liquid nitrogen for DNA extraction. For bacterial abundance analysis, we filled 5-ml cryovials (VWR) with 2.5-3 grams of sediment containing a 10% solution of paraformaldehyde/glutaraldehyde^69^ in 0.22μm-filtered streamwater that we added *in-situ*, and we flash froze the vials in liquid nitrogen.

### Measurement of *in-situ* physicochemical parameters and chlorophyll-α content

We measured stream temperature, dissolved oxygen and pH using a WTW Multi-parameter portable meter (MultiLine® Multi 3630 IDS), conductivity using a WTW - IDS probe (TetraCon® 925) and turbidity using a WTW portable turbidity meter (Turb® 430 IR). We measured the concentration of chlorophyll *a* in the sediment following a modified ethanol extraction protocol as described elsewhere^70^. Geographical and physicochemical parameters are shown in Table S5.

### Bacterial abundance

We quantified the number of cells per gram of dry sediment using flow cytometry after detaching the cells from the sediment matrix, by slightly modifying the method of Amalfitano & Fazi^71^ as described elsewhere^70^. Briefly, we fixed 2.5 - 3 grams of wet sediment per sample *in situ* in 1.8 ml of filter-sterilized paraformaldehyde/glutaraldehyde solution^69^ within cryovials and we flash-froze the vials in liquid nitrogen. At the end of the expedition, we transferred the samples in dry ice back to the lab and we stored them at -80°C until the analysis. To detach the cells, we performed two rounds of mild shaking (Standard Analog Shaker, VWR, 15 min, 5.5 speed) followed by sonication (Sonifier 450, Branson, 1 min, 60% duty cycle, output 5) in 10 ml of paraformaldehyde/glutaraldehyde solution supplemented with sodium pyrophosphate at a final concentration of 0.025 mM. We pooled the resulting extracts per sample (∼20-22 ml in total), mixed them thoroughly, transferred 1 ml of each in a sterile 1.5 ml tube and spinned them for 5 sec to pellet the large sediment particles. We then diluted 100 µl of the supernatants 10-fold in paraformaldehyde/glutaraldehyde solution and we stained the dilutions with SybrGreen® (1X final concentration, incubation for 15 min at 37°C) before analyzing them on a NovoCyte flow cytometer (ACEA Biosciences) equipped with a 488 nm laser.

To analyze the stained samples we set the reading time to 2 min per sample and the flow rate to 14 µl per min, rinsing thrice and shaking once between samples. We identified and gated the cell populations based on the height of their fluorescence signals on a 530/30 – 725/40 nm biplot as previously described^72^ (Fig. S5), using the ACEA NovoExpress® software with thresholds of 300 and 3000 on the front scatter and 530/30 nm channels, respectively. We analyzed three stained technical replicates plus one unstained replicate of the same extract per sample, the latter to exclude any background fluorescence. The coefficient of variation among the counts from technical replicates was 7.5±5.1% on average. Finally, we corrected the acquired numbers for the various dilution factors and for the sediments’ water content (which we obtained from the weight loss of oven-dried sediment samples) to obtain the amount of total cells per gram of dry sediment.

### DNA extraction, PCR amplification and 16S rRNA gene amplicon sequencing

We extracted DNA from sediment samples using a phenol-chloroform method with certain modifications to address the nature of our samples^73^.We amplified the V3-V4 hypervariable regions of the bacterial 16S rRNA gene using primers 341f (5’-CCTACGGGNGGCWGCAG-3’) and 785r (5’-GACTACHVGGGTATCTAATCC-3’)^74^. Due to low DNA yields and presence of inhibitors in the DNA extracts of certain samples and in an attempt to avoid PCR biases due to unequal input DNA, we diluted all DNA samples to a final concentration of ≤ 2-3 ng/ul. The KAPA HiFi DNA Polymerase (Hot Start and Ready Mix formulation) was used in a 25-ul-amplification reaction containing 1X PCR buffer, 1 uM of each primer, 0.48 ug/ul BSA and 1.0 ul of template DNA. Amplification was performed in a Biometra Trio (Biometra) instrument. The thermal conditions applied after an initial denaturation at 95°C for 3 min, were 94°C for 30 s, 55°C for 30 s and 72°C for 30 s for 25 cycles followed by a final extension at 72°C for 5 min. Amplification was verified on a 1.5% agarose gel and products were sent to Lausanne Genomic Technologies Facility (Switzerland) for further processing, library preparation and 300-base-pairs paired-end sequencing at an Illumina Miseq platform.

### Sequence downstream analyses

We used Trimmomatic v.0.36^75^ for quality filtering of the sequencing reads. Shortly, we truncated the reads in 4bp sliding windows at the first instance of mean quality dropping below a Phred score of 15, we removed the three leading and trailing nucleotides and we discarded the reads that were shorter than 200bp.

We performed all subsequent sequence processing within the QIIME2 v.2019.1 framework^76^. We used DADA2^77^ with the default parameters to remove the primers, denoise and join the reads into exact sequence variants (SVs). For this, 17 and 21 nucleotides (corresponding to the primers’ length) were removed at the beginning of the forward and reverse reads respectively, and the reads were truncated at 300bp. We performed denoising and joining of the reads using the default parameters, and we removed any SVs that were not found in at least two samples. We used the *alpha-rarefaction* method implemented in the *diversity* plugin of QIIME2 to create the rarefaction curves (Fig. S6). We used the SV table that contained the raw sequence counts of each SV at each sample to calculate the relative abundances of SVs within samples, and we transformed the relative abundances into absolute abundances (cells per gram of dry sediment) by multiplying with the cell counts derived from flow cytometry^78^.

We assigned taxonomy with the *feature-classifier* plugin^79^ in QIIME2. First, we trained QIIME2’s naïve Bayesian classifier using the *fit-classifier-naïve-bayes* method on the Greengenes^80^ 99% OTUS database v. 13.5. We created this training set using the *extract-reads* method with a minimal and maximal length of 250 and 550 nucleotides, respectively, and using the primers’ sequences. Finally, we assigned the taxonomy of the sequence variants using the *classify-sklearn* method with default parameters. We considered the taxonomies down to the genus level, ignoring “species” assignments that can be ambiguous based only on part of the 16S rRNA gene^81^. At the Class level, Betaproteobacteria were dominant in all samples (Fig. S7). A detailed taxonomic summary can be found in Supplementary Results.

To build the phylogenetic tree, we aligned the sequences of the SVs with *mafft*^82^ and we trimmed the alignment with the *mask* method in QIIME2 using the default parameters. We then used RAxML^83^ with the GTRCAT substitution model and the rapid bootstrap option to build the tree, and the *midpoint-root* method to root the phylogenetic tree. To calculate pairwise nucleotidic similarities we used ClustalOmega^84^ v.1.2.3.

### Identification of the core microbiome

We identified the core microbiome across all samples based on taxonomy, i.e., as the consensus taxonomic clades that are present in all 40 reaches (20 GFS x 2 reaches each). We used the package *metacoder*^85^ in R^86^ to visualize the results as hierarchy trees.

### Multivariate statistics

We used distance-based redundancy analysis to quantify the variance in the Bray-Curtis similarity matrix (calculated using log-transformed absolute abundances) that could be explained by the measured physicochemical variables, using the *capscale()* function of the *vegan*^87^ package in R. We performed a stepwise forward selection based on the increase in the adjusted R^2^ to select for the variables to include in the model, using the *ordiR2step()* function in *vegan* with 200 permutations (Table S6).

### Quantification of the dominant assembly processes at the community level

We used the framework developed by Stegen and colleagues^20,21^ to quantify the dominant assembly processes at the community level. This framework assigns differences between two given communities to selection (either homogeneous or heterogeneous), to dispersal (either homogenizing or limiting) or to the lack of any dominant process. The influence of selection is first determined by examining the phylogenetic β-diversity via the z-score (in this case called β-nearest taxon index - βNTI) of the observed β-mean nearest taxon distance (β-MNTD) from a null distribution of the same metric. βNTI scores less than -2 indicate that the observed β-MNTD is significantly smaller than ∼95% of the null values and thus that homogeneous selection between the compared communities causes them to have more similar species phylogenetically (at short distances) than expected by chance. In analogy, βNTI scores greater than +2 indicate the dominance of heterogeneous selection. Community pairs with βNTI scores between -2 and +2 are further compared in terms of composition using the Raup-Crick distances based on the Bray-Curtis similarity (RC_Bray_), with the null distribution in this case being formed by probabilistic permutations under weak selection and random dispersal. Here, values of RC_Bray_ less than -0.95 and greater than 0.95 indicate less and more compositional turnover, respectively, than the null expectation and that is attributed to homogenizing dispersal in the former case and to dispersal limitation in the latter.

To apply the framework, two main assumptions must hold true for the examined dataset: some degree of migration occurred among local communities at least at some point in evolutionary time and b) phylogenetic conservatism exists, i.e., phylogenetically more similar organisms occupy more similar ecological niches. For our dataset, the first assumption probably holds true even for the most distant GFS because at some point in geological time all the sampled locations were covered under the same ice sheet. To test the second assumption of phylogenetic conservatism we first calculated the niche optima of the SVs for each physicochemical parameter that we included in the multivariate analyses (Table S6), as previously described^88^, and we then calculated the niche distances among SVs as the euclidean distance of their niche optima (after standardization of each parameter). We then performed a mantel correlogram analysis, correlating the phylogenetic distances to the niche distances at different distance classes. Proper use of the βMNTD requires a positive correlation between the two at short genetic distance classes, indicating that at short phylogenetic distances more related SVs have shorter niche optima distances and therefore occupy more similar ecological niches; that was indeed the case for our dataset (Figure S2). We calculated the abundance-weighted βMNTD using the *comdistnt* function of the *picante*^89^ package in R (setting abundance.weighted = TRUE).

### Identification of phylogenetic clades under homogeneous selection

In addition to inferring community-wise patterns of assembly using the framework of Stegen et al. as described above, we further developed a framework to identify phylogenetic clades under homogeneous selection. In analogy to the community-wise framework, we defined those clades as phylogenetically coherent units containing SVs with phylogenetically closer relatives across communities than expected by chance.

Our method consists of the following steps:

1. For a given pair of communities and for each SV therein that is not present in both communities, we calculate its z-score from a null distribution of log-transformed minimum phylogenetic distances to assess how different its minimum phylogenetic distance is to that expected by chance. For example, if we examine SV *x* in the community pair *a-b* and *x* is present in community *a* and not in community *b*, we first find the minimum phylogenetic distance *dist* between *x* and all the SVs that are present in community *b* and we log-transform it. We then calculate a null distribution of minimum log-transformed phylogenetic distances between *x* and these SVs by randomly picking phylogenetic distances from the whole metacommunity distance pool and assigning them to the SVs in the two communities, with *null_mean* and *null_sd* being the mean and standard deviation of this distribution. Here, we performed 100 permutations to calculate *null_mean* and *null_sd*, because this number was adequate to create normally distributed null distances. For future studies we recommend that null distributions are checked for normality depending on the sample size of each study and that the number of permutations is adjusted accordingly. Finally, we calculate the z-score as:

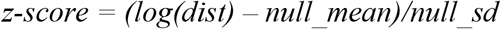
2. We then calculate for each SV its total z-score across all community pairs. We use phylofactorization^29,30^ to identify phylogenetically coherent groups of SVs with significantly different total scores compared to outgroups and to extract the consensus taxonomic classification of the SVs within.
3. We also detect SVs with constantly low phylogenetic turnover (LPT-SVs). For that, we calculate the median z-score and we classify LPT-SVs using a threshold of significance based on a median z-score less than -2. We perceive this threshold as an indication that a given SV is consistently under homogeneous selection, because this SV has phylogenetically nearest neighboring SVs with shorter distances than those expected by chance in at least 50% of the applicable community pairs (i.e., those community pairs where the SV is found in exactly one of the two communities). While this classification was tailored for the purposes of our study, the concept can be expanded in future studies to potentially detect and classify SVs based on other numerical threshold or even SVs with higher phylogenetic turnover than that expected by chance. In the latter case the numerical threshold should be positive and it depends on the study to find appropriate criteria. If, for instance, heterogeneous selection is detected at the community level for a subset of community pairs, a threshold of +2 regarding the median could be used to detect SVs with consistently high phylogenetic turnover among the majority of these community pairs.

## Supporting information

Supplementary Results, Supplementary Figures and Supplementary Tables

## Acknowledgements

This work was supported by NOMIS Foundation under the “Vanishing Glaciers” project granted to TJB. The funders had no role in study design, data collection and analysis, decision to publish, or preparation of the manuscript.

## Author contributions

TJB, HP and SF conceived and designed the study; MSt, MT, VDS and HP performed sampling; PP, HP, TK, JB, MSch, SB and SF performed lab work; SF and MB analyzed the sequencing data; AW and SF developed new code; TJB and PW provided resources; SF, TJB and HP wrote the manuscript with input from all authors.

## Data Availability

Sequencing data has been uploaded to the European Nucleotide Archive under accession number PRJEB40567.

## Code Availability

The R code for the calculation of the phylogenetic z-scores has been uploaded to GitHub (https://github.com/sfodel/phylo_z_scores).

## Conflict of Interest Statement

The authors declare that they have no conflict of interest

## Notes

https://github.com/sfodel/phylo_z_scores

### Competing Interest Statement

The authors have declared no competing interest.

## References

1 Larkin, A. A. & Martiny, A. C. Microdiversity shapes the traits, niche space, and biogeography of microbial taxa. Environ. Microbiol. Rep. 9, 55–70 (2017).

2 Acinas, S. G. et al. Fine-scale phylogenetic architecture of a complex bacterial community. Nature 430, 551–554 (2004).

3 Thompson, J. R. et al. Genotypic diversity within a natural coastal bacterioplankton population. Science 307, 1311 (2005).

4 Thrash, C. J. et al. Single-cell enabled comparative genomics of a deep ocean SAR11 bathytype. ISME J. 8, 1440–1451 (2014).

5 Hunt, D. E. et al. Resource partitioning and sympatric differentiation among closely related bacterioplankton. Science 320, 1081 (2008).

6 Kent, A. G. et al. Parallel phylogeography of Prochlorococcus and Synechococcus. ISME J. 13, 430–441 (2019).

7 Brown, M. V. & Furham, J. A. Marine bacterial microdiversity as revealed by internal transcribed spacer analysis. Aquat. Microb. Ecol. 41, 15–23 (2005).

8 Scanlan, D. J. et al. Ecological genomics of marine picocyanobacteria. Microbiol. Mol. Biol. Rev. 73, 249 (2009).

9 Yung, C.-M. et al. Thermally adaptive tradeoffs in closely related marine bacterial strains. Environ. Microbiol. 17, 2421–2429 (2015).

10 Props, R. & Denef, V. J. Temperature and nutrient levels correspond with lineage-specific microdiversification in the ubiquitous and abundant freshwater genus Limnohabitans. Appl. Environ. Microbiol. 86, e00140–00120 (2020).

11 Chase, A. B. et al. Microdiversity of an abundant terrestrial bacterium encompasses extensive variation in ecologically relevant traits. mBio 8 (2017).

12 Choudoir, M. J. & Buckley, D. H. Phylogenetic conservatism of thermal traits explains dispersal limitation and genomic differentiation of Streptomyces sister-taxa. ISME J. 12, 2176–2186 (2018).

13 Martiny, J. B., Jones, S. E., Lennon, J. T. & Martiny, A. C. Microbiomes in light of traits: A phylogenetic perspective. Science 350, 9323 (2015).

14 Vellend, B. M. Conceptual synthesis in community ecology. Quart. Rev. Biol. 85, 183–206 (2010).

15 Chafee, M. et al. Recurrent patterns of microdiversity in a temperate coastal marine environment. ISME J. 12, 237–252 (2018).

16 Needham, D. M., Sachdeva, R. & Fuhrman, J. A. Ecological dynamics and co-occurrence among marine phytoplankton, bacteria and myoviruses shows microdiversity matters. ISME J. 11, 1614–1629 (2017).

17 Garcia-Garcia, N., Tamames, J., Linz, A. M., Pedros-Alio, C. & Puente-Sanchez, F. Microdiversity ensures the maintenance of functional microbial communities under changing environmental conditions. ISME J. 13, 2969–2983 (2019).

18 Tromas, N. et al. The evolution of realized niches within freshwater Synechococcus. Environ. Microbiol. 22, 1238–1250 (2020).

19 Konstantinidis, K. T., Ramette, A. & Tiedje, J. M. The bacterial species definition in the genomic era. Philos. Trans. R. Soc. B: Biol. Sci. 361, 1929–1940 (2006).

20 Stegen, J. C. et al. Quantifying community assembly processes and identifying features that impose them. ISME J. 7, 2069–2079 (2013).

21 Stegen, J. C., Lin, X., Fredrickson, J. K. & Konopka, A. E. Estimating and mapping ecological processes influencing microbial community assembly. Front. Microbiol. 6 (2015).

22 Allen, R., Hoffmann, L. J., Larcombe, M. J., Louisson, Z. & Summerfield, T. C. Homogeneous environmental selection dominates microbial community assembly in the oligotrophic South Pacific Gyre. Mol. Ecol. 29, 4680–4691, (2020).

23 Li, Y. et al. Homogeneous selection dominates the microbial community assembly in the sediment of the Three Gorges Reservoir. Sci. Tot. Environ. 690, 50–60 (2019).

24 Zhang, K. et al. Salinity is a key determinant for soil microbial communities in a desert ecosystem. mSystems 4 (2019).

25 Cauvy-Fraunié, S. & Dangles, O. A global synthesis of biodiversity responses to glacier retreat. Nat. Ecol. Evol. 3, 1675–1685 (2019).

26 Wilhelm, L., Singer, G. A., Fasching, C., Battin, T. J. & Besemer, K. Microbial biodiversity in glacier-fed streams. ISME J. 7, 1651 (2013).

27 Milner, A. M. et al. Glacier shrinkage driving global changes in downstream systems. Proc. Nat. Acad. Sci. USA 114, 9770 (2017).

28 Fodelianakis, S. et al. Dispersal homogenizes communities via immigration even at low rates in a simplified synthetic bacterial metacommunity. Nat. Commun. 10, 1314, (2019).

29 Washburne, A. D. et al. Phylogenetic factorization of compositional data yields lineage-level associations in microbiome datasets. PeerJ 5, e2969 (2017).

30 Washburne, A. D. et al. Phylofactorization: a graph partitioning algorithm to identify phylogenetic scales of ecological data. Ecol. Monogr. 89, e01353, (2019).

31 Logares, R. et al. Biogeography of bacterial communities exposed to progressive long-term environmental change. ISME J. 7, 937–948 (2013).

32 Cerqueira, T., Barroso, C., Froufe, H., Egas, C. & Bettencourt, R. Metagenomic signatures of microbial communities in deep-sea hydrothermal sediments of Azores Vent Fields. Microb. Ecol. 76, 387–403 (2018).

33 Osburn, M. R., LaRowe, D. E., Momper, L. M. & Amend, J. P. Chemolithotrophy in the continental deep subsurface: Sanford underground research facility (SURF), USA. Front. Microbiol. 5 (2014).

34 Tran, P. et al. Microbial life under ice: Metagenome diversity and in situ activity of Verrucomicrobia in seasonally ice-covered Lakes. Environ. Microbiol. 20, 2568–2584 (2018).

35 Vick-Majors, T. J., Priscu, J. C. & Amaral-Zettler, L. A. Modular community structure suggests metabolic plasticity during the transition to polar night in ice-covered Antarctic lakes. ISME J. 8, 778–789 (2014).

36 Darcy, J. L., Lynch, R. C., King, A. J., Robeson, M. S. & Schmidt, S. K. Global distribution of Polaromonas phylotypes -evidence for a highly successful dispersal capacity. PloS One 6, e23742 (2011).

37 Smith, H. J., Foreman, C. M. & Ramaraj, T. Draft genome sequence of a metabolically diverse Antarctic supraglacial stream organism, Polaromonas sp. strain CG9_12, determined using Pacific Biosciences single-molecule real-time sequencing technology. Genome Announc. 2, e01242–01214 (2014).

38 Gawor, J. et al. Evidence of adaptation, niche separation and microevolution within the genus Polaromonas on Arctic and Antarctic glacial surfaces. Extremophiles 20, 403–413 (2016).

39 Rime, T., Hartmann, M. & Frey, B. Potential sources of microbial colonizers in an initial soil ecosystem after retreat of an alpine glacier. ISME J. 10, 1625–1641 (2016).

40 Liu, Q., Zhou, Y.-G. & Xin, Y.-H. High diversity and distinctive community structure of bacteria on glaciers in China revealed by 454 pyrosequencing. Syst. Appl. Microbiol. 38, 578–585 (2015).

41 Kalyuzhnaya, M. G., Bowerman, S., Lara, J. C., Lidstrom, M. E. & Chistoserdova, L. Methylotenera mobilis gen. nov., sp. nov., an obligately methylamine-utilizing bacterium within the family Methylophilaceae. Int. J. Syst. Evol. Microbiol. 56, 2819–2823 (2006).

42 Kane, S. R. et al. Whole-genome analysis of the methyl tert-butyl ether-degrading Beta-Proteobacterium Methylibium petroleiphilum PM1. J. Bacteriol. 189, 1931, (2007).

43 Martineau, C., Mauffrey, F., Villemur, R. & Müller, V. Comparative analysis of denitrifying activities of Hyphomicrobium nitrativorans, Hyphomicrobium denitrificans, and Hyphomicrobium zavarzinii. Appl. Environ. Microbiol. 81, 5003–5014 (2015).

44 Dieser, M. et al. Molecular and biogeochemical evidence for methane cycling beneath the western margin of the Greenland Ice Sheet. ISME J. 8, 2305–2316 (2014).

45 Michaud, A. B. et al. Microbial oxidation as a methane sink beneath the West Antarctic Ice Sheet. Nat. Geosci. 10, 582–586 (2017).

46 Baker, J. M. et al. Genome sequence of Rhodoferax antarcticus ANT.BRT; a psychrophilic purple nonsulfur bacterium from an Antarctic microbial mat. Microorganisms 5, (2017).

47 Crisafi, F., Giuliano, L., Yakimov, M. M., Azzaro, M. & Denaro, R. Isolation and degradation potential of a cold-adapted oil/PAH-degrading marine bacterial consortium from Kongsfjorden (Arctic region). Rendiconti Lincei 27, 261–270 (2016).

48 Zhong, Z.-P. et al. Clean low-biomass procedures and their application to ancient ice core microorganisms. Front. Microbiol. 9 (2018).

49 Bai, Y. et al. Variation in denitrifying bacterial communities along a primary succession in the Hailuogou Glacier retreat area, China. PeerJ 7:e7356 (2019).

50 Garcia-Lopez, E., Rodriguez-Lorente, I., Alcazar, P. & Cid, C. Microbial communities in coastal glaciers and tidewater tongues of Svalbard archipelago, Norway. Front. Mar. Sci. 5 (2019).

51 Liu, S. et al. Comammox Nitrospira within the Yangtze River continuum: community, biogeography, and ecological drivers. ISME J. 14, 2488–2504 (2020).

52 Harrold, Z. R. et al. Aerobic and anaerobic thiosulfate oxidation by a cold-adapted, subglacial chemoautotroph. Appl. Environ. Microbiol. 82, 1486–1495 (2016).

53 Franzetti, A. et al. Early ecological succession patterns of bacterial, fungal and plant communities along a chronosequence in a recently deglaciated area of the Italian Alps. FEMS Microbiol. Ecol. 96 (2020).

54 Kohler, T. J., Van Horn, D. J., Darling, J. P., Takacs-Vesbach, C. D. & McKnight, D. M. Nutrient treatments alter microbial mat colonization in two glacial meltwater streams from the McMurdo Dry Valleys, Antarctica. FEMS Microbiol. Ecol. 92:4 (2016).

55 Sawayama, M. et al. Isolation of a Leptothrix strain, OUMS1, from ocherous deposits in groundwater. Cur. Microbiol. 63, 173–180 (2011).

56 Li, Y. et al. Reconstruction of the functional ecosystem in the high light, low temperature union glacier region, Antarctica. Front. Microbiol. 10 (2019).

57 Milner, A. M. et al. Glacier shrinkage driving global changes in downstream systems. Proc. Nat. Acad. Sci. USA 114, 9770–9778 (2017).

58 Jorquera, M. A., Graether, S. P. & Maruyama, F. Editorial: bioprospecting and biotechnology of extremophiles. Front. Bioeng. Biotech. 7, 204 (2019).

59 Stegen, J. C., Lin, X., Konopka, A. E. & Fredrickson, J. K. Stochastic and deterministic assembly processes in subsurface microbial communities. ISME J. 6, 1653–1664 (2012).

60 Zhou, J. & Ning, D. Stochastic community assembly: does it matter in microbial ecology? Microbiol. Mol. Biol. Rev. 81(4) (2017).

61 Ning, D., Deng, Y., Tiedje, J. M. & Zhou, J. A general framework for quantitatively assessing ecological stochasticity. Proc. Nat. Acad. Sci. USA 116, 16892–16898 (2019).

62 Zhou, J. et al. Stochasticity, succession, and environmental perturbations in a fluidic ecosystem. Proc. Nat. Acad. Sci. USA 111, E836–845 (2014).

63 Ning, D. et al. A quantitative framework reveals ecological drivers of grassland microbial community assembly in response to warming. Nat. Commun. 11, 4717 (2020).

64 Leibold, M. A. et al. The metacommunity concept: a framework for multi-scale community ecology. Ecol. Let. 7, 601–613 (2004).

65 Chase, A. B. et al. Emergence of soil bacterial ecotypes along a climate gradient. Environ. Microbiol. 20, 4112–4126 (2018).

66 Callahan, B. J., Grinevich, D., Thakur, S., Balamotis, M. A. & Yehezkel, T. B. Ultra-accurate microbial amplicon sequencing rirectly from complex samples with synthetic long reads. bioRxiv, (2020).

67 Matsuo, Y. et al. Full-length 16S rRNA gene amplicon analysis of human gut microbiota using MinION™ nanopore sequencing confers species-level resolution. bioRxiv (2020).

68 Nygaard, A. B., Tunsjø, H. S., Meisal, R. & Charnock, C. A preliminary study on the potential of Nanopore MinION and Illumina MiSeq 16S rRNA gene sequencing to characterize building-dust microbiomes. Sci. Rep. 10, 3209 (2020).

69 Duarte, C. M. et al. Discovery of Afifi, the shallowest and southernmost brine pool reported in the Red Sea. Sci. Rep. 10, 910 (2020).

70 Kohler, T. J. et al. Patterns and drivers of extracellular enzyme activity in New Zealand glacier-fed streams. Front. Microbiol. 11, 2922 (2020).

71 Amalfitano, S. & Fazi, S. Recovery and quantification of bacterial cells associated with streambed sediments. J. Microbiol. Methods 75, 237–243 (2008).

72 Hammes, F. et al. Flow-cytometric total bacterial cell counts as a descriptive microbiological parameter for drinking water treatment processes. Water Res. 42, 269–277 (2008).

73 Busi, S. B. et al. Optimised biomolecular extraction for metagenomic analysis of microbial biofilms from high-mountain streams. PeerJ 8, e9973, (2020).

74 Klindworth, A. et al. Evaluation of general 16S ribosomal RNA gene PCR primers for classical and next-generation sequencing-based diversity studies. Nucleic Acids Res. 41(1), e1 (2013).

75 Bolger, A. M., Lohse, M. & Usadel, B. Trimmomatic: a flexible trimmer for Illumina sequence data. Bioinformatics 30, 2114–2120 (2014).

76 Bolyen, E. et al. Reproducible, interactive, scalable and extensible microbiome data science using QIIME 2. Nat. Biotech. 37, 852–857 (2019).

77 Callahan, B. J. et al. DADA2: High-resolution sample inference from Illumina amplicon data. Nat. Meth. 13, 581–583 (2016).

78 Props, R. et al. Absolute quantification of microbial taxon abundances. ISME J. 11, 584–587 (2017).

79 Bokulich, N. A. et al. Optimizing taxonomic classification of marker-gene amplicon sequences with QIIME 2’s q2-feature-classifier plugin. Microbiome 6, 90 (2018).

80 DeSantis, T. Z. et al. Greengenes, a chimera-checked 16S rRNA gene database and workbench compatible with ARB. Appl. Environ. Microbiol. 72, 5069–5072 (2006).

81 Singer, E. et al. High-resolution phylogenetic microbial community profiling. ISME J. 10, 2020–2032 (2016).

82 Katoh, K. & Standley, D. M. MAFFT multiple sequence alignment software version 7: improvements in performance and usability. Mol. Biol. Evol. 30, 772–780 (2013).

83 Stamatakis, A. RAxML version 8: a tool for phylogenetic analysis and post-analysis of large phylogenies. Bioinformatics 30, 1312–1313 (2014).

84 Sievers, F. et al. Fast, scalable generation of high-quality protein multiple sequence alignments using Clustal Omega. Mol. Syst. Biol. 7, 539–539 (2011).

85 Foster, Z. S. L., Sharpton, T. J. & Grünwald, N. J. Metacoder: An R package for visualization and manipulation of community taxonomic diversity data. PLOS Comput. Biol. 13, e1005404 (2017).

86 R: A Language and Environment for Statistical Computing (R Foundation for Statistical Computing, Vienna, Austria, 2014).

87 Oksanen, J., et al. The vegan package. Community ecology package 10.631-637 (2007): 719. (2013).

88 Fodelianakis, S. et al. Modified niche optima and breadths explain the historical contingency of bacterial community responses to eutrophication in coastal sediments. Mol Ecol. 26, 2006–2018 (2017).

89 Kembel, S. W. et al. Picante: R tools for integrating phylogenies and ecology. Bioinformatics 26, 1463–1464 (2010).

