## Supplementary Results, Supplementary Figures and Supplementary Tables for "Homogeneous selection promotes microdiversity in the glacier-fed stream microbiome"

### Detailed taxonomic diversity

The communities in the different glacier-fed streams were both rich in terms of the number of detected SVs and in terms of the number of taxonomic groups. We detected 8186 SVs in total and 640 ± 60 SVs per glacier-fed stream on average. In terms of taxonomy, there were 36 assigned bacterial Phyla, 112 Classes, 162 Orders, 156 Families and 159 Genera present in the sampled communities. The most diverse and abundant Phyla were Proteobacteria (3267 SVs, 34.4-90.2% of total cells per sample), Bacteroidetes (1324 SVs, 1.8-31.9% of total cells per sample) and Planctomycetes (728 SVs, 0.5-9.9% of total cells per sample) and the most diverse and abundant Classes within these three phyla were Betaproteobacteria (1429 SVs, 15.2-78.8% of total cells per sample), Saprospirae (481 SVs, 0.09-13.4% of total cells per sample) and Planctomycetia (539 SVs, 0.4-8.5% of total cells per sample), respectively (Fig. S6).

### Core microbiome

The core microbiome, which we defined as the taxonomic units present in at least one sample at every reach, included 11 Phyla, 20 Classes, 29 Orders, 18 Families and 11 Genera (Fig. 1, Table S2). The 11 Genera within the core microbiome included a total of 1133 SVs (13.9% of the total SVs) that comprised on average 34.8% (13.6 – 62%) of the total cells per gram of dry sediment per community. Six core Genera were taxonomically affiliated to Betaproteobacteria; *Methylotenera, Polaromonas, Rhodoferax, Leptothrix, Methylibium* and *Rubrivivax,* containing 541 SVs and comprising on average 25.1% of the total cells per gram of dry sediment per community. Two core Genera were affiliated to Planctomycetes; *Gemmata* and

*Planctomyces*, containing 258 SVs and comprising on average 1.3% of the total cells per gram of dry sediment per community. Two core Genera were affiliated to Alphaproteobacteria; *Hyphomicrobium* and *Novosphingobium*, containing 113 SVs and comprising on average 3.5% of the total cells per gram of dry sediment per community. One core Genus was affiliated to Bacteroidetes; *Flavobacterium*, containing 205 SVs and comprising on average 3% of the total cells per gram of dry sediment per community, and one Genus was affiliated to Nitrospirae; *Nitrospira*, containing 20 SVs and comprising on average 1.9% of the total cells per gram of dry sediment per community.

**Environmental drivers of bacterial β-diversity**

Using distance-based redundancy analysis and a forward stepwise model building, we found that seven of the recorded environmental parameters, namely conductivity, dissolved oxygen, latitude, chl-α content, pH, water temperature and turbidity, could collectively explain 43.5% of the variance in the Bray-Curtis (BC) dissimilarity among the samples (Fig. S4, Table S6). Conductivity had the highest proportion of variance explained (14.1%), followed by dissolved oxygen (8.1%), latitude (5.7%), chl-α content (5.6%) and pH (4.3%) while the rest of the variables explained low proportions of variance (1.4-2.3%).

The samples had a distinct clustering pattern along the first constrained axis that explained 18.8% of the total BC variance (Fig. S4). The left axis side was characterized by a low-dispersion cluster of samples from turbulent streams with high conductivity, pH and dissolved oxygen, and the right axis side was characterized by a high-dispersion cluster of samples from warmer streams with higher chl-α content. The second constrained axis explained 8.7% of total BC variance and was mostly associated with latitude, suggesting that a part of community variability can be explained by geographic legacies. Samples from the same stream but at different reaches (UP and DN) clustered mostly together and the variance explained by the factor “reach” was non-significant.

### Supplementary Figures

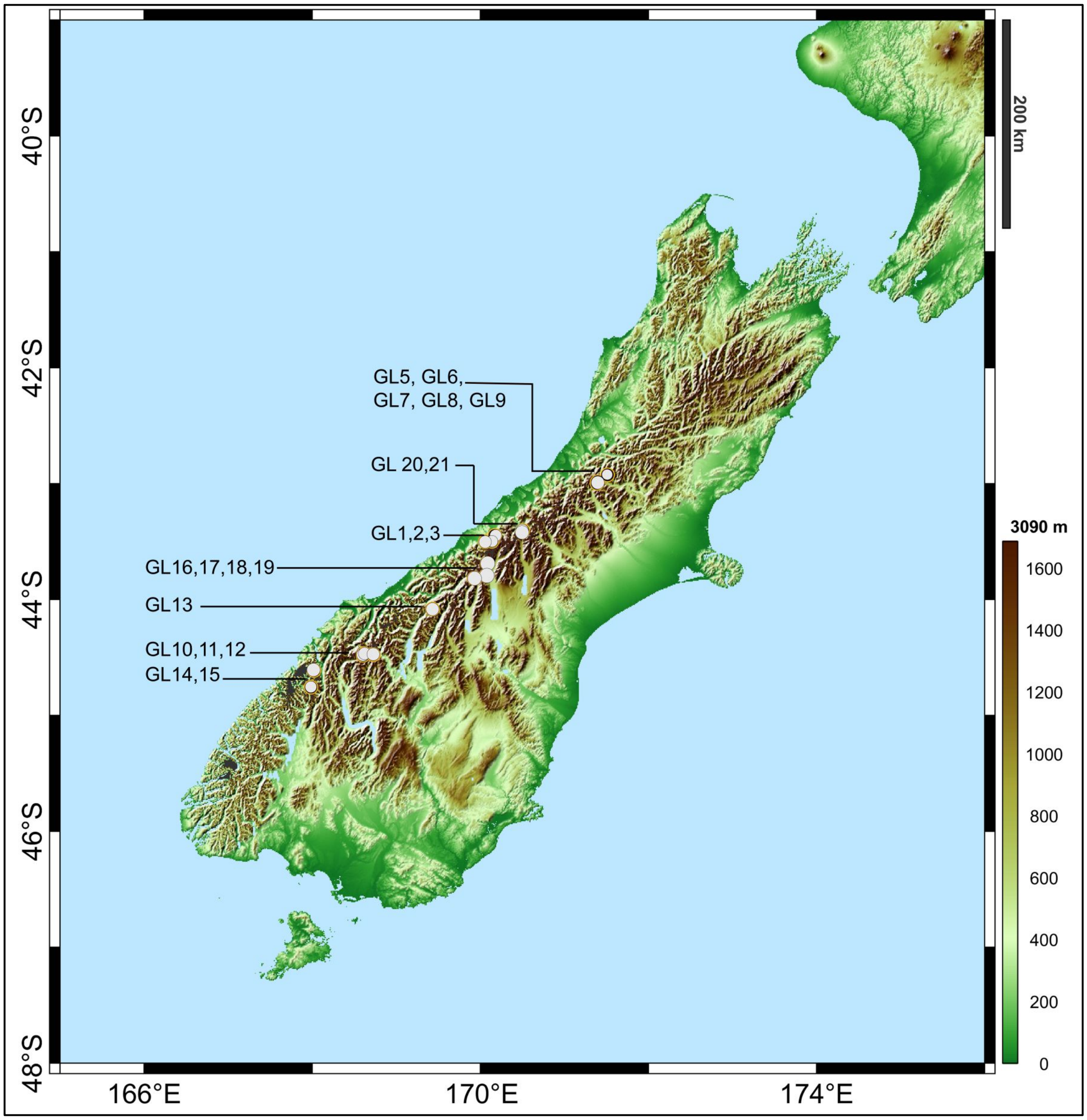


**Figure S1. The location of the sampled glacier-fed streams (GFS) at the Southern Alps of New Zealand.** GFS are assigned numbers from 1 to 21 (excluding 4) for operational purposes. Map colors correspond to the altitude as per the legend on the right.

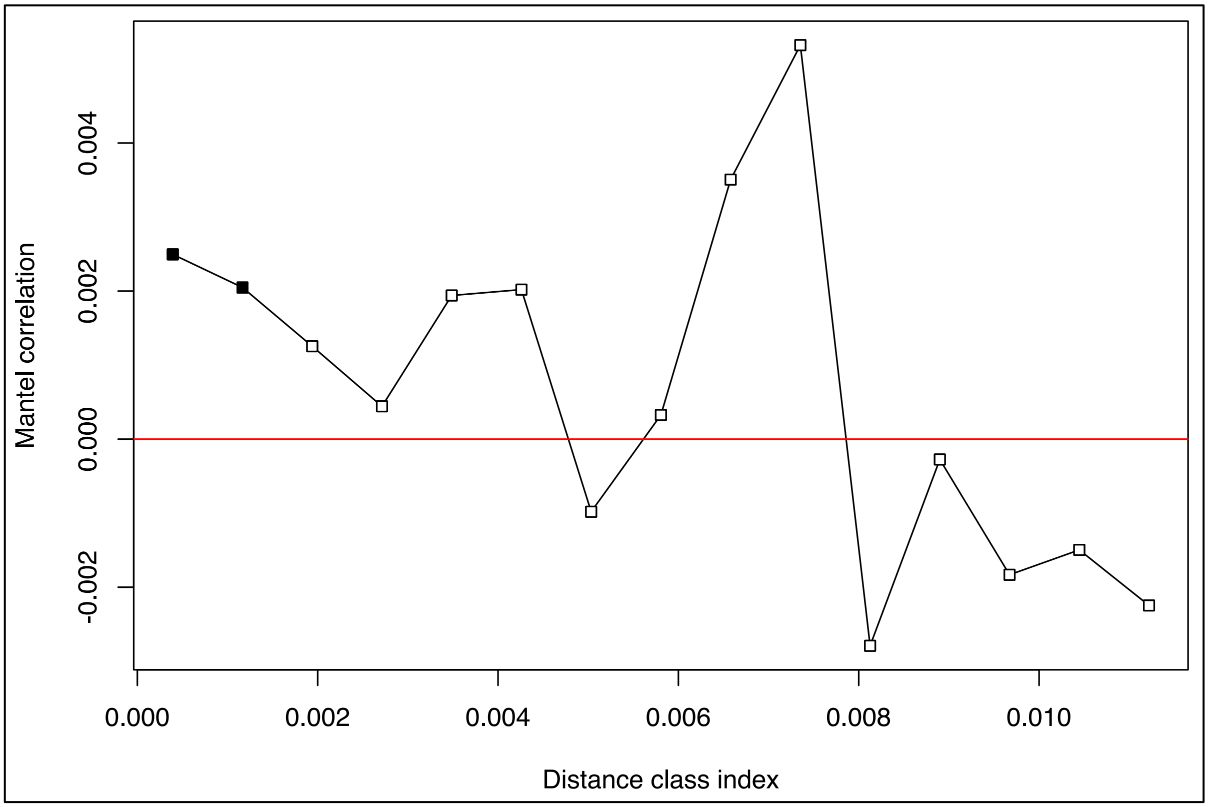


**Figure S2. Mantel correlogram between phylogenetic distances and niche distances among all the Sequence Variants.** Filled squares correspond to significant correlations and empty squares correspond to non-significant correlations. The phylogenetic distances were Hellinger-transformed before the correlations. The niche distances were calculated as the Euclidean distances of standardized niche optima regarding the environmental parameters that explained a significant proportion of β-diversity as per the db-RDA analysis, namely conductivity, dissolved oxygen, latitude, chl-α, pH, water temperature and turbidity.

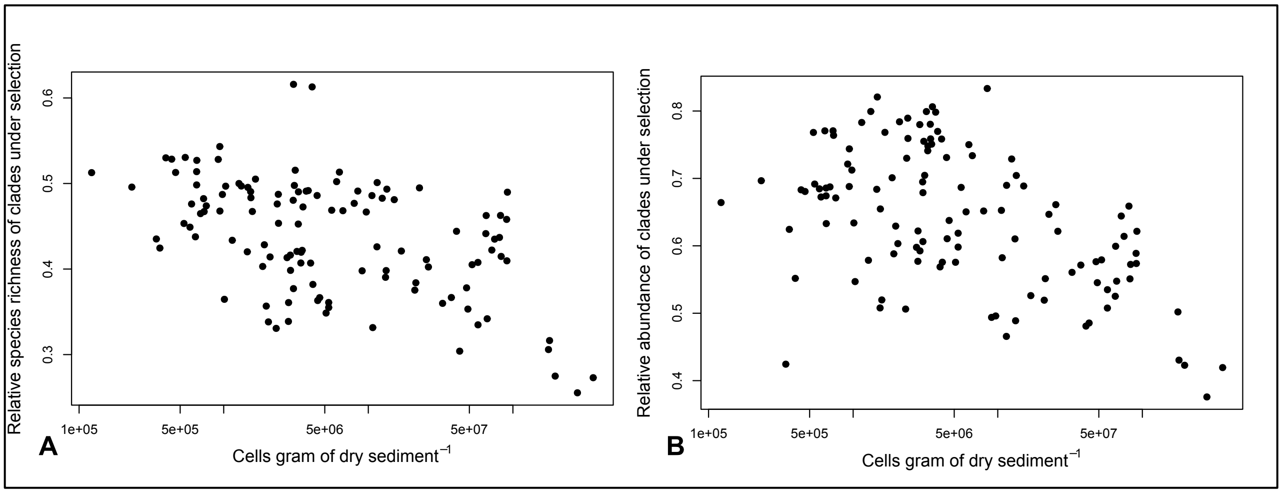


**Figure S3. The inverse relationship between the relative species richness (A) and the relative abundance (B) of phylogenetic clades under environmental selection and total bacterial cell density A. The relative species richness of the clades under environmental selection (y-axis) as a function of total bacterial cell density (x-axis, in log-scale). B. The relative abundance of the clades under environmental selection (y-axis) as a function of total bacterial cell density (x-axis, in log-scale).** Adjusted $R^2$= 0.252, and 0.282, for panels A and B, respectively. For both panels $p<<0.001$ and $n=119$.

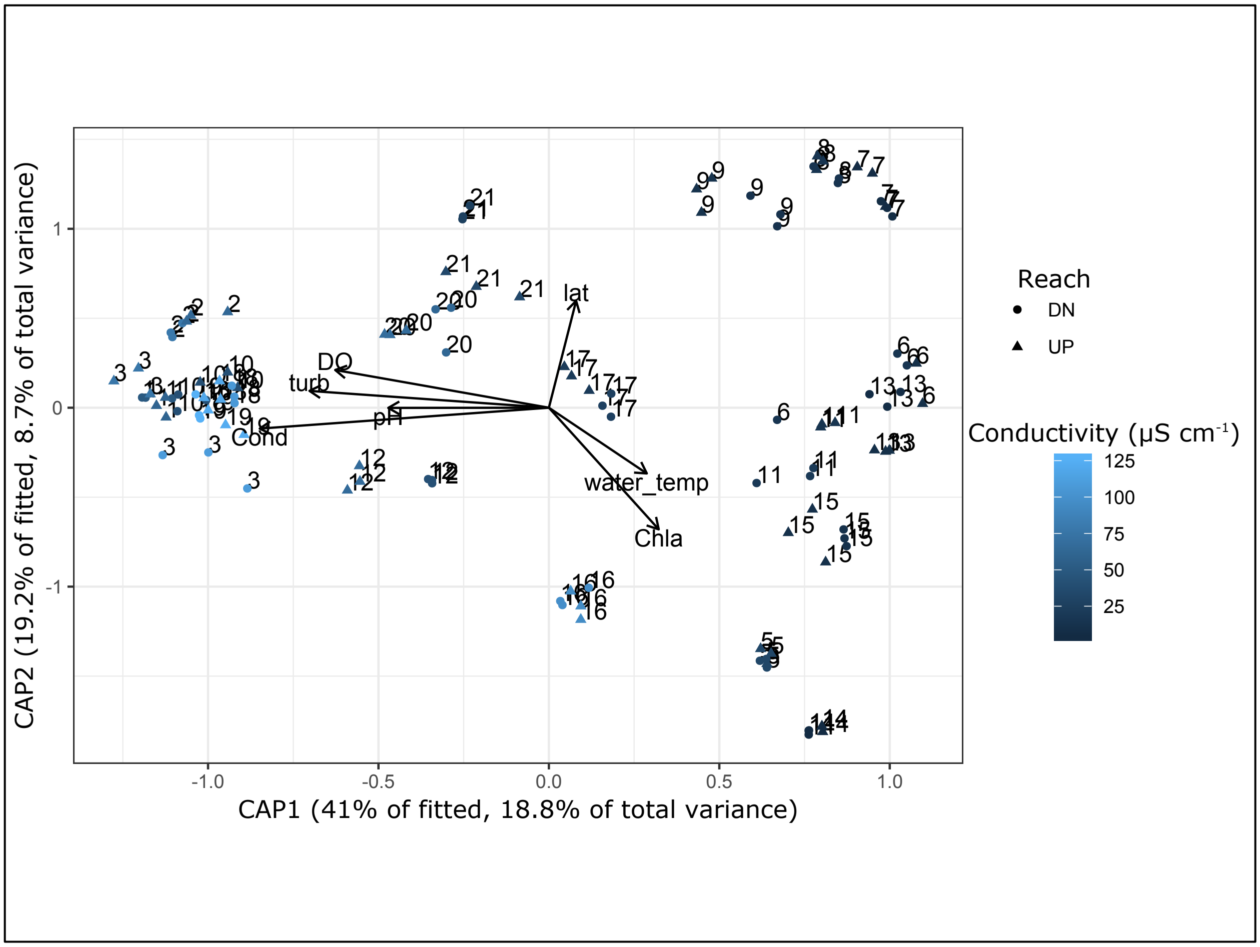


**Figure S4. The constrained ordination plot of the distance-based redundancy analysis on the BC dissimilarity of bacterial communities from all the sampled streams.** Factors included in the model are plotted as vectors. Samples are colored based on conductivity that was the factor explaining most of the BC variance, as per the legend on the middle right. Sample symbols are given according to the reach as per the legend on the upper right. Numbers above samples indicate the location of the stream as per Figure S1. Axes are scaled based on the variance they explain. Cond: conductivity, DO: dissolved oxygen, lat: latitude, Chla: chl-α content, water_temp: water temperature, turb: turbidity.

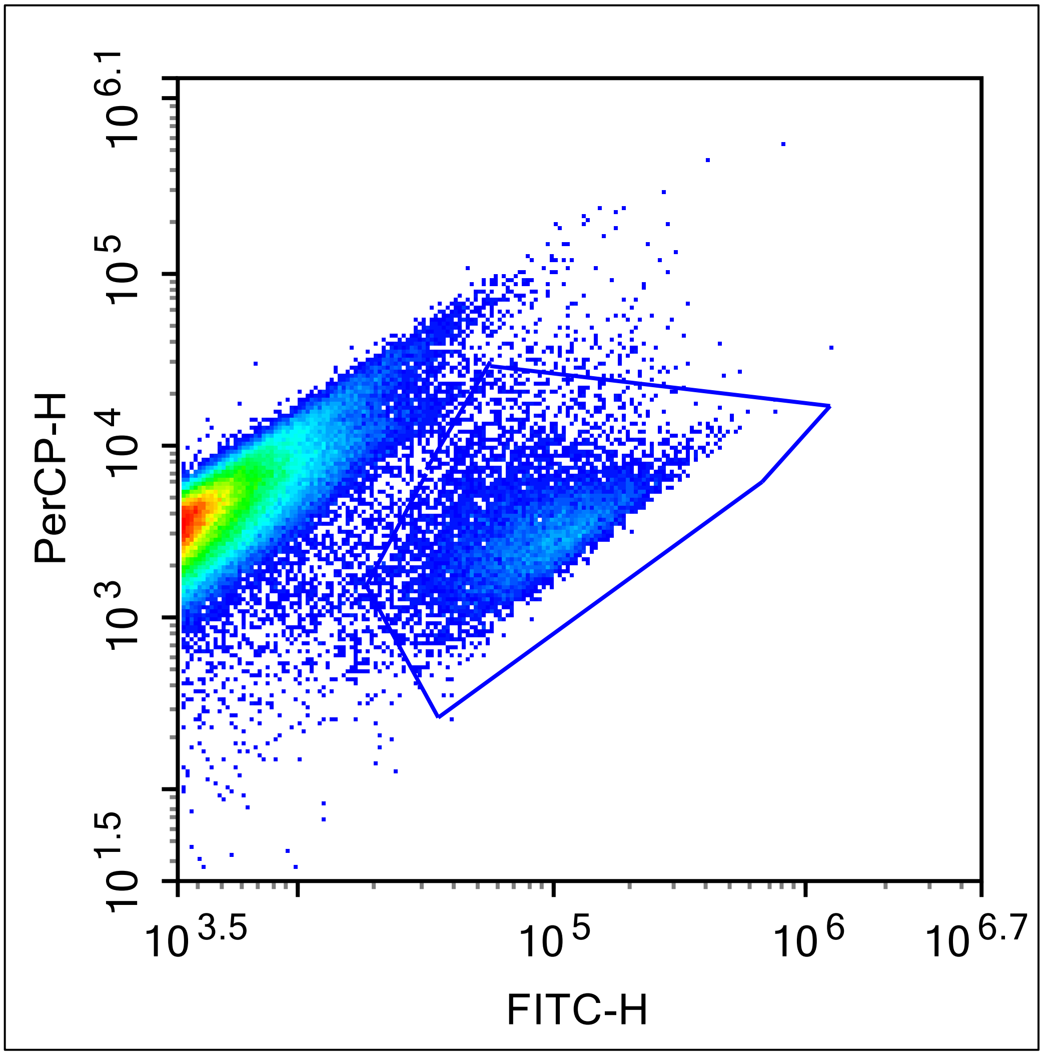


**Figure S5. Flow cytometry scatterplot of green (FITC-H – x-axis) versus red (PerCP-H – y-axis) fluorescence of a typical sediment fixed extract.** The polygon delineates the gate used for counting cells, which is based on events with disproportionately more green fluorescence compared to red fluorescence because of SybrGreen staining. On the contrary, debris is shown on the middle left as a dense cluster of events with higher red:green fluorescence ratio.

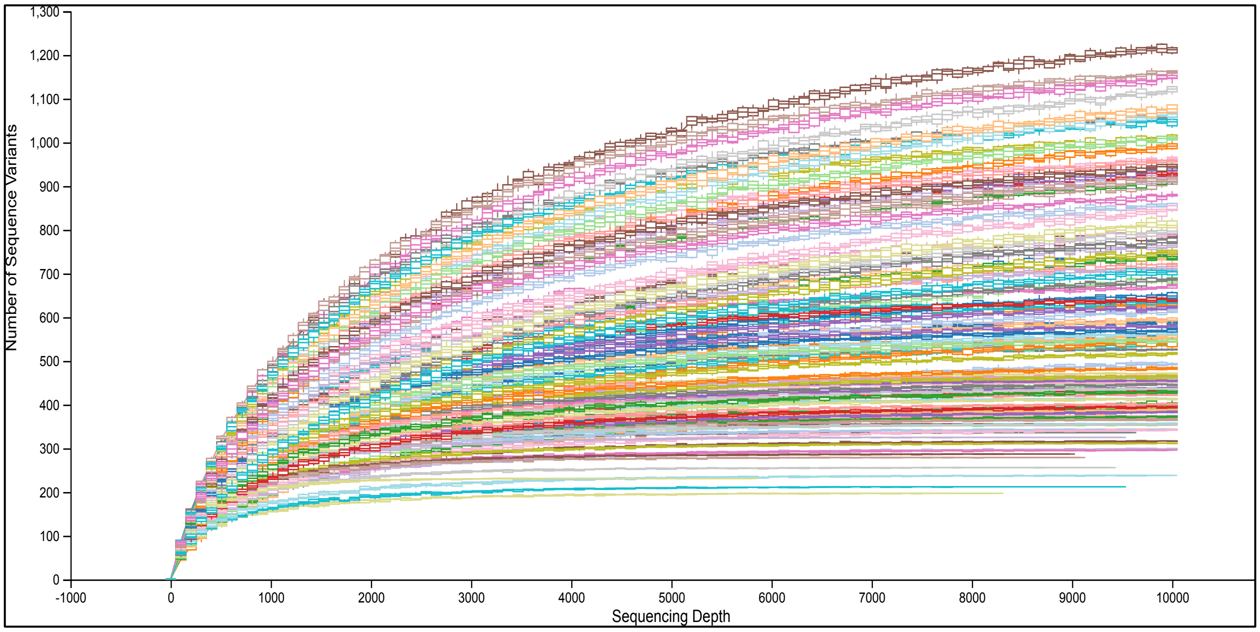


**Figure S6. The rarefaction curves of the generated 16S rRNA gene amplicons from all the samples.** The number of sequences is shown on the x-axis and the number of observed sequence variants (SVs) are shown on the y-axis. Rarefaction was performed at intervals of 100 sequences with 10 permutations per sequence interval at a sequencing depth of up to 10,000 reads. The range of the obtained values is represented with boxplots. Different colors represent different samples.

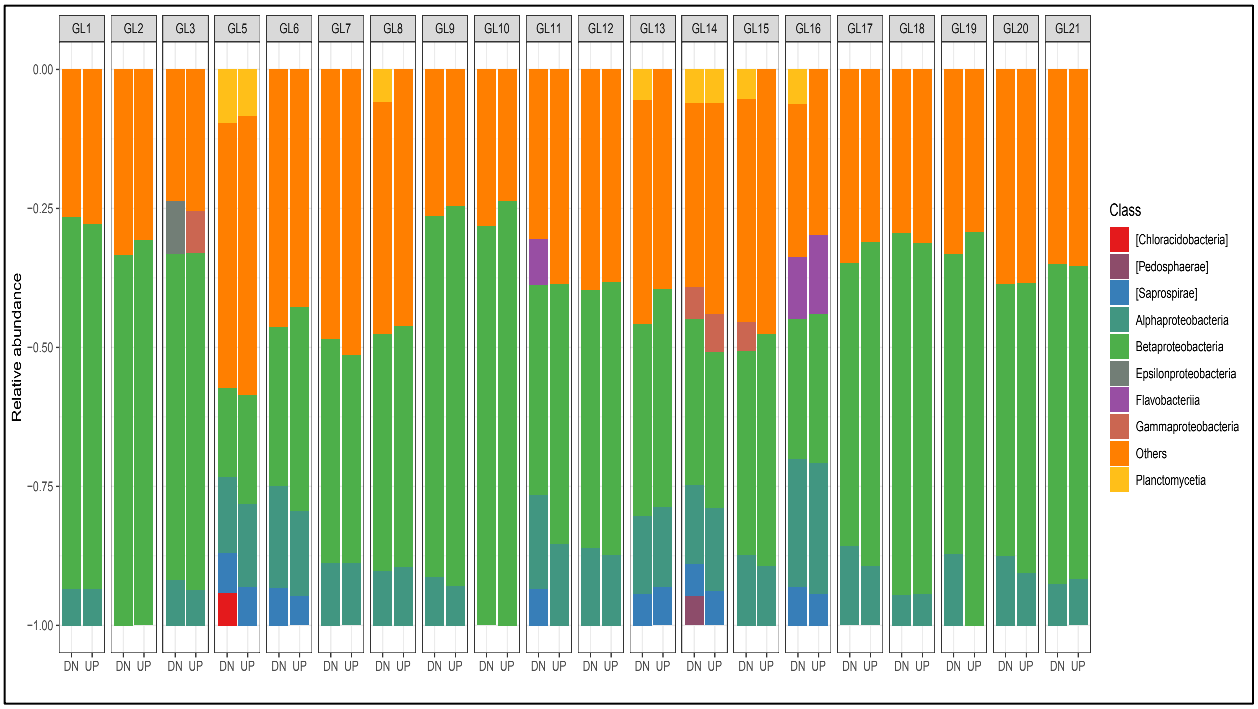


**Figure S7. Taxonomical composition of the sampled communities at the Class level.** The sampled streams are ordered by ascending operational number. Each barplot corresponds to a different reach (UP or DN) within a given stream and represents the mean of three (or two in the case of GL6 – UP) within-reach samples. Classes with low mean relative abundances are grouped together in the "Others" category. Candidate classes are named in brackets.

### Supplementary Tables

**Table S1. The identified phylogenetic groups under homogeneous selection, i.e., with significantly lower total z-scores compared to outgroups.** Group 1 corresponds to the phylogenetic group having different scores and Group 2 to the phylogenetic outgroup that the comparison is made against.

| Factor | Group 1 Consensus Taxonomy | Number of SVs in Group 1 | Group 2 Consensus Taxonomy | Number of SVs in Group 2 | Contrast test p-value |
|---|---|---|---|---|---|
| **1** | Betaproteobacteria (Class) | 1418 | All the rest | 6758 | 6.8E-255 |
| **2** | Novosphingobium (Genus) | 5 | All present Bacteria except Betaproteobacteria | 6753 | 8.5E-148 |
| **3** | Nitrospira (Genus) | 18 | All present Bacteria except Betaproteobacteria | 6731 | 3.4E-67 |
| **4** | Alphaproteobacteria (Class) | 602 | All present Bacteria except Betaproteobacteria | 6129 | 6.4E-70 |
| **5** | [Saprospirae] (Candidate Class) | 338 | All present Bacteria except Alpha- and Betaproteobacteria | 5791 | 3.3E-59 |
| **6** | Methylotenera (Genus) | 48 | Betaproteobacteria | 1359 | 2.3E-17 |
| **7** | Comamonadaceae (Family) | 575 | Betaproteobacteria except Methylotenera | 784 | 3.3E-16 |
| **8** | Ellin606 (Uncultured Order) | 54 | Betaproteobacteria except Methylotenera and Comamonadaceae | 730 | 2.5E-18 |

**Table S2. The core microbiome, i.e., the taxonomic units present in at least one sample at every reach, at different taxonomic levels.**

| Core Level | Number of Taxa | Taxa names | Number of SVs within the taxa | Mean relative abundance (min-max) |
|---|---|---|---|---|
| **Phylum** | 11 | Proteobacteria, Nitrospirae, Bacteroidetes, Planctomycetes, Acidobacteria, Actinobacteria, OD1, Verrucomicrobia, Chloroflexi, Gemmatimonadetes, Cyanobacteria | 7554 | 94.3% (78-97.1%) |
| **Class** | 20 | Chloracidobacteria, Acidobacteria-6, Solibacteres, Acidimicrobia, Actinobacteria, Thermophilia, Saprospirae, Cytophagia, Flavobacteriia, Sphingobacteriia, Chloroplast, Gemmatimonadetes, Nitrospira, ZB2, Planctomycetia, Alphaproteobacteria, Betaproteobacteria, Gammaproteobacteria, Deltaproteobacteria, Spartobacteria | 6103 | 91% (77.1-96.3%) |
| **Order** | 29 | Rhizobiales, Sphingomonadales, Rhodobacterales, Caulobacterales, Pseudomonadales, MND1, Ellin6067, IS-44, KD8-87, Acidimicrobiales, Solibacterales, iii1-15, Gemmatales, Sphingobacteriales, RB41, Gaiellales, Chthoniobacterales, Burkholderiales, Methylophilales, Actinomycetales, Flavobacteriales, Cytophagales, Saprospirales, Planctomycetales, Pirellulales, Nitrospirales, Myxococales, Xanthomonadales | 4514 | 72.5% (30.4-91%) |
| **Family** | 18 | Hyphomicrobiaceae, Sphingomonadaceae, Caulobacteraceae, Chthoniobacteraceae, Saprospiraceae, Ellin6075, Gemmataceae, Sinobacteraceae, Oxalobacteraceae, Comamonadaceae, Methylophilaceae, Flavobacteriaceae, Cytophagaceae, Chitinophagaceae, Planctomycetaceae, Pirellulaceae, Nitrospiraceae, Xanthomonadaceae | 2724 | 49.2% (19.8-72.2%) |
| **Genus** | 12 | Methylotenera, Polaromonas, Rhodoferax, Leptothrix, Methylibium, Rubrivivax, Novosphingobium, Hyphomicrobium, Nitrospira, Flavobacterium, Planctomyces, Gemmata | 1133 | 34.8% (13.6-62%) |

**Table S3. The properties of the detected genera across the whole dataset that had at least 2 Sequence Variants (SVs) and that are formally assigned in the scientific nomenclature.**

| | Number of SVs | Mean Nucleotidic Similarity (%)[#] | Mean Niche Breadth (B) | Mean relative abundance |
|---|---|---|---|---|
| **Flavobacterium** | 201 | 93,02 | 3,91 | 2,95E-02 |
| **Planctomyces** | 153 | 86,46 | 4,18 | 1,00E-02 |
| ***Methylibium** | 130 | 96,59 | 4,62 | 1,58E-02 |
| ***Rhodoferax** | 112 | 96,66 | 4,93 | 2,87E-02 |
| **Gemmata** | 105 | 86,13 | 3,56 | 3,27E-03 |
| ***Leptothrix** | 98 | 96,45 | 4,98 | 2,46E-02 |
| ***Methylotenera** | 84 | 95,42 | 6,99 | 1,24E-01 |
| **Bdellovibrio** | 84 | 85,46 | 2,73 | 1,91E-03 |
| ***Hyphomicrobium** | 82 | 95,16 | 5,24 | 1,27E-02 |
| ***Polaromonas** | 73 | 96,55 | 6,93 | 4,94E-02 |
| ***Rhodobacter** | 69 | 95,93 | 5,50 | 1,30E-02 |
| ***Thiobacillus** | 48 | 95,09 | 7,37 | 4,67E-02 |
| ***Rubrivivax** | 44 | 95,81 | 4,78 | 8,82E-03 |
| **DA101** | 44 | 93,90 | 3,33 | 2,14E-03 |
| **Hymenobacter** | 36 | 94,58 | 2,35 | 8,57E-04 |
| ***Novosphingobium** | 31 | 96,58 | 6,07 | 2,20E-02 |
| ***Janthinobacterium** | 26 | 97,37 | 2,15 | 7,56E-04 |
| ***Herminiimonas** | 24 | 96,12 | 3,11 | 1,85E-03 |
| ***Rhodoplanes** | 22 | 95,64 | 3,38 | 1,09E-03 |
| ***Kaistobacter** | 22 | 97,18 | 6,02 | 6,91E-03 |
| ***Sediminibacterium** | 20 | 94,51 | 5,79 | 2,86E-03 |
| ***Nitrospira** | 20 | 96,53 | 8,12 | 1,90E-02 |
| **Luteolibacter** | 20 | 97,16 | 2,81 | 1,09E-03 |
| **Opitutus** | 19 | 95,94 | 2,71 | 3,67E-04 |
| **Candidatus Solibacter** | 17 | 94,26 | 3,68 | 8,83E-04 |
| **Gemmatimonas** | 15 | 96,70 | 3,51 | 4,27E-04 |
| **CM44** | 15 | 94,13 | 3,71 | 6,40E-04 |
| **Deinococcus** | 15 | 91,93 | 2,55 | 2,87E-04 |
| ***Segetibacter** | 12 | 93,87 | 5,92 | 1,29E-03 |
| **A17** | 12 | 93,74 | 3,51 | 5,13E-04 |
| **Candidatus Koribacter** | 11 | 92,67 | 3,22 | 2,66E-04 |
| **Fimbriimonas** | 11 | 90,08 | 4,07 | 5,37E-04 |
| ***Phenylobacterium** | 10 | 95,78 | 4,50 | 9,00E-04 |
| ***Paucibacter** | 10 | 98,26 | 4,09 | 5,69E-04 |
| **Chthoniobacter** | 10 | 96,86 | 4,22 | 8,80E-04 |
| ***Devosia** | 9 | 96,47 | 3,18 | 3,35E-04 |
| **Geobacter** | 9 | 95,96 | 2,67 | 2,94E-04 |

| | | | | |
|---|---|---|---|---|
| **Phormidium** | 8 | 96,55 | 2,91 | 2,52E-04 |
| **Pirellula** | 8 | 90,23 | 3,46 | 1,39E-04 |
| ***Gallionella** | 8 | 95,25 | 7,60 | 4,79E-03 |
| **Pseudomonas** | 8 | 97,96 | 3,03 | 4,78E-04 |
| **Lysobacter** | 8 | 96,40 | 3,90 | 5,53E-04 |
| **Rudanella** | 7 | 92,11 | 2,00 | 5,34E-05 |
| ***Flavisolibacter** | 7 | 94,70 | 2,95 | 1,35E-04 |
| ***Sphingomonas** | 7 | 96,06 | 3,68 | 3,95E-04 |
| ***Ramlibacter** | 7 | 95,45 | 3,11 | 1,97E-04 |
| ***Variovorax** | 7 | 96,45 | 4,30 | 9,50E-04 |
| **Leptospira** | 7 | 92,54 | 3,86 | 1,69E-04 |
| **Cytophaga** | 6 | 92,75 | 3,00 | 1,28E-04 |
| **Spirosoma** | 6 | 89,42 | 2,03 | 2,94E-05 |
| **Pseudanabaena** | 6 | 91,41 | 3,14 | 1,84E-04 |
| ***Hydrogenophaga** | 6 | 96,94 | 1,42 | 8,03E-04 |
| ***Dechloromonas** | 6 | 97,49 | 1,76 | 1,63E-04 |
| **Cellvibrio** | 6 | 95,96 | 3,31 | 1,28E-04 |
| **Aquicella** | 6 | 90,38 | 2,38 | 4,06E-05 |
| **Salinibacterium** | 5 | 98,16 | 3,78 | 3,15E-04 |
| **Fluviicola** | 5 | 92,65 | 2,53 | 3,60E-05 |
| **JG37-AG-70** | 5 | 98,23 | 3,27 | 1,62E-04 |
| ***Pedomicrobium** | 5 | 97,96 | 3,18 | 8,82E-05 |
| ***Zymomonas** | 5 | 98,33 | 2,53 | 6,96E-05 |
| **Sulfuricurvum** | 5 | 95,28 | 4,68 | 1,45E-02 |
| **Leadbetterella** | 4 | 97,68 | 4,71 | 3,51E-04 |
| **Sporocytophaga** | 4 | 91,96 | 1,92 | 2,14E-05 |
| **Leptolyngbya** | 4 | 88,99 | 2,09 | 6,27E-05 |
| ***Pelomonas** | 4 | 97,19 | 7,72 | 1,47E-03 |
| ***Sulfuritalea** | 4 | 96,45 | 3,71 | 7,73E-05 |
| **Crenothrix** | 4 | 94,48 | 4,33 | 7,69E-05 |
| **Bryobacter** | 3 | 97,51 | 8,08 | 3,67E-04 |
| **Armatimonas** | 3 | 99,34 | 5,38 | 7,13E-04 |
| **Paludibacter** | 3 | 96,68 | 2,63 | 5,25E-05 |
| **Emticicia** | 3 | 95,90 | 3,56 | 1,02E-04 |
| **Candidatus Protochlamydia** | 3 | 93,99 | 2,58 | 1,94E-05 |
| **Candidatus Rhabdochlamydia** | 3 | 88,29 | 1,93 | 5,18E-06 |
| **Gloeobacter** | 3 | 85,83 | 1,84 | 2,59E-05 |
| **Calothrix** | 3 | 96,86 | 2,11 | 6,80E-05 |
| ***Bosea** | 3 | 97,60 | 6,13 | 4,07E-04 |
| ***Labrys** | 3 | 97,18 | 2,25 | 1,22E-05 |
| ***Lutibacterium** | 3 | 96,93 | 2,36 | 4,99E-05 |
| **Smithella** | 3 | 96,50 | 3,12 | 9,75E-06 |
| **Arcobacter** | 3 | 98,84 | 1,67 | 7,38E-04 |

| | | | | |
|---|---|---|---|---|
| **Gluconacetobacter** | 3 | 98,75 | 5,77 | 1,84E-04 |
| **Acinetobacter** | 3 | 96,34 | 4,30 | 9,39E-05 |
| **Steroidobacter** | 3 | 94,54 | 1,87 | 1,07E-05 |
| **Iamia** | 2 | 98,76 | 1,36 | 8,01E-06 |
| **Demequina** | 2 | 97,54 | 1,60 | 1,07E-05 |
| **Crocinitomix** | 2 | 97,87 | 5,78 | 5,96E-05 |
| ***Niabella** | 2 | 99,05 | 2,59 | 1,60E-05 |
| **Haliscomenobacter** | 2 | 99,29 | 1,46 | 1,33E-05 |
| **Dolichospermum** | 2 | 99,01 | 1,81 | 2,31E-05 |
| **Staphylococcus** | 2 | 98,83 | 1,83 | 3,85E-06 |
| **Clostridium** | 2 | 94,53 | 1,57 | 8,06E-06 |
| **BD2-6** | 2 | 90,65 | 2,35 | 8,13E-06 |
| ***Bradyrhizobium** | 2 | 99,75 | 4,24 | 1,86E-04 |
| ***Methylobacterium** | 2 | 96,77 | 3,70 | 4,83E-05 |
| ***Rubellimicrobium** | 2 | 92,79 | 2,32 | 9,49E-06 |
| ***Phaeospirillum** | 2 | 97,51 | 6,08 | 2,92E-05 |
| ***Sphingobium** | 2 | 96,02 | 1,57 | 4,32E-05 |
| ***Delftia** | 2 | 98,59 | 1,84 | 9,04E-06 |
| **Plesiocystis** | 2 | 99,76 | 2,87 | 1,25E-05 |
| **HTCC** | 2 | 99,53 | 1,51 | 2,57E-06 |
| **Thiofaba** | 2 | 98,13 | 2,37 | 6,81E-06 |
| **Legionella** | 2 | 96,49 | 2,18 | 1,00E-05 |
| **Alkanindiges** | 2 | 99,53 | 2,09 | 5,59E-05 |
| **Perlucidibaca** | 2 | 92,04 | 3,20 | 7,09E-06 |
| **Rhodanobacter** | 2 | 98,36 | 1,64 | 1,98E-05 |
| **Thermomonas** | 2 | 98,59 | 3,45 | 4,70E-05 |
| **Prosthecobacter** | 2 | 96,01 | 3,77 | 3,15E-05 |
| **Pedosphaera** | 2 | 93,90 | 2,03 | 1,48E-05 |
| **Candidatus Xiphinematobacter** | 2 | 90,62 | 2,34 | 4,17E-06 |
| **OR-59** | 2 | 81,67 | 3,41 | 8,81E-06 |

*Genus within phylogenetic clades under homogeneous selection [#]Based on the sequenced part of the 16S rRNA gene

**Table S4. The properties of the Sequence Variants with constantly low phylogenetic turnover (LPT-SVs).** LPT-SVs are grouped based on their consensus taxonomy and groups are sorted based on the percent of total score (column 3).

| Consensus Taxonomy (Level) | Number of LPT-SVs | Percent of Total Score among LPT-SVs* | Mean Total Score per LPT-SV | Mean contributing community pairs per LPT-SV |
|---|---|---|---|---|
| *Nitrospira (genus) | 16 | 13.1 | -6481 | 1866 |
| *Methylotenera (genus) | 29 | 12 | -3262 | 1352 |
| *Polaromonas (genus) | 20 | 11.3 | -4455 | 1515 |
| *Ellin6067 (order) | 10 | 10.9 | -8615 | 2147 |
| *Leptothrix (genus) | 19 | 8.8 | -3647 | 1241 |
| *Burkholderiales (order) | 13 | 8.8 | -5301 | 1412 |
| *Novosphingobium (genus) | 8 | 6.8 | -6751 | 1310 |
| Pirellulaceae (family) | 3 | 4.6 | -11992 | 2560 |
| mb2424 (family) | 6 | 2.7 | -3562 | 1119 |
| Sphingobacteriales (order) | 3 | 2.7 | -7071 | 1915 |
| *Betaproteobacteria (class) | 9 | 2.7 | -2441 | 913 |
| *Rhodoferax (genus) | 5 | 2.3 | -3601 | 994 |
| *Thiobacillus (genus) | 3 | 2 | -5135 | 1231 |
| Rhizobiales (order) | 2 | 1.9 | -7423 | 2319 |
| Ellin6075 (family) | 2 | 1.8 | -7013 | 3482 |
| Gaiellaceae (family) | 1 | 1.4 | -10893 | 2051 |
| Xanthomonadaceae (family) | 4 | 1.2 | -2385 | 573 |
| Chitinophagaceae (family) | 2 | 0.8 | -3619 | 1100 |
| Chromatiales (order) | 1 | 0.7 | -5457 | 3074 |
| *Rhodobacter (genus) | 1 | 0.6 | -4272 | 880 |
| *Saprospiraceae (family) | 1 | 0.5 | -3849 | 2346 |
| *MND1 (order) | 2 | 0.4 | -1573 | 971 |
| *Methylibium (genus) | 3 | 0.4 | -983 | 492 |
| *IS-44 (order) | 3 | 0.3 | -896 | 419 |
| Flectobacillus (genus) | 1 | 0.3 | -2444 | 456 |
| Planctomyces (genus) | 1 | 0.2 | -1621 | 348 |
| Sulfuricurvum (genus) | 1 | 0.2 | -1592 | 342 |
| Bacteroidetes (phylum) | 1 | 0.2 | -1261 | 566 |
| *Methylophilaceae (family) | 1 | 0.2 | -1229 | 674 |
| Gammaproteobacteria (class) | 1 | 0.1 | -1060 | 230 |
| **Total for *** | **143** | **80.5** | | |

* Groups residing within phylogenetic clades under homogeneous selection

**Table S5. The measured geographical and physicochemical properties at each sampled glacier-fed stream.**

| Glacier | Reach | Altitude (m.a.s.l.) | Water T (°C) | DO (mg $l^{-1}$) | pH | Potential (mV) | Conductivity (μS $cm^{-1}$) | Turbidity (NTU) | Chl-α (μg $g^{-1}$) |
|---|---|---|---|---|---|---|---|---|---|
| GL1 | UP | 354 | 2.4 | 13.3 | 7.81 | -47.7 | 60.1 | 363 | 1.02E-04 |
| GL1 | DN | 210 | 3.1 | 13.4 | 7.71 | -42.9 | 49.4 | 347 | 9.93E-05 |
| GL2 | UP | 1186 | 0 | 12.8 | 9.11 | 119.4 | 64 | 227 | 2.3E-04 |
| GL2 | DN | 1100 | 1.23 | 12.75 | 8.32 | n/d | 78.8 | 164.33 | 2.42E-04 |
| GL3 | UP | 266 | 0.33 | 15.38 | 8.83 | -104 | 77.2 | 265.33 | 3.63E-04 |
| GL3 | DN | 219 | 4.3 | 12.72 | 8.31 | -77.3 | 109.2 | 352.67 | 6.84E-04 |
| GL5 | UP | 1326 | 7.7 | 10.35 | 7.54 | -34.1 | 32.4 | 0.05 | 1.58E-01 |
| GL5 | DN | 1264 | 9.35 | 9.89 | 7.55 | -34.9 | 32.5 | 0 | 2.58E-01 |
| GL6 | UP | 1362 | 8.03 | 10.31 | 7.6 | -39.3 | 9.2 | 7.82 | 1.04E-02 |
| GL6 | DN | 1117 | 7.95 | 10.49 | 8.15 | -72.8 | 15 | 0.92 | 2.95E-03 |
| GL7 | UP | 1779 | 3 | 11.04 | 6.72 | 11.5 | 5.9 | 37.7 | 7.48E-04 |
| GL7 | DN | 1707 | 3.58 | 10.98 | 6.84 | 5.5 | 6.3 | 32.33 | 7.7E-04 |
| GL8 | UP | 1670 | 0.68 | 11.86 | 7.34 | -22.4 | 14.9 | 12.57 | 6.18E-04 |
| GL8 | DN | 1604 | 1.25 | 11.74 | 7.17 | -12.7 | 14.2 | 11.1 | 7.37E-04 |
| GL9 | UP | 1391 | 3.58 | 11.51 | 7.19 | -12.8 | 12.5 | 23.07 | 5.14E-04 |
| GL9 | DN | 1246 | 6.03 | 10.99 | 6.78 | 8.7 | 12.2 | 27.23 | 5.5E-04 |
| GL10 | UP | 1111 | 0.65 | 13.16 | 8.1 | -64 | 40.7 | 48.97 | 7.51E-04 |
| GL10 | DN | 1077 | 0.7 | 12.92 | 7.43 | -27.2 | 40.8 | 98.57 | 8.14E-04 |
| GL11 | UP | 1581 | 2.93 | 11.14 | 10.18 | -180 | 14.5 | 4.12 | 1.1E-02 |
| GL11 | DN | 1474 | 5.38 | 10.65 | 10.18 | -181.6 | 19 | 1.92 | 2.67E-02 |
| GL12 | UP | 756 | 6.6 | 11.35 | 9.83 | -163.1 | 66.7 | 9.88 | 3.33E-03 |
| GL12 | DN | 714 | 7 | 11.39 | 9.83 | -163.1 | 41.7 | 8.22 | 3.4E-03 |
| GL13 | UP | 1720 | 0.73 | 11.58 | 6.76 | 15.2 | 9.25 | 4.95 | 1.78E-02 |
| GL13 | DN | 1662 | 0.85 | 11.93 | 6.7 | 19 | 6.9 | 4.53 | 8.84E-03 |
| GL14 | UP | 1079 | 3.38 | 11.81 | 6.41 | 34.3 | 4.5 | 0 | 1.151E-01 |
| GL14 | DN | 1056 | 4.85 | 11.37 | 6.23 | 44.5 | 4.5 | 0 | 7.17E-02 |
| GL15 | UP | 1281 | 1.4 | 11.94 | 6.47 | 30.9 | 10.4 | 0.81 | 3.85E-03 |
| GL15 | DN | 1245 | 1.6 | 12.04 | 6.6 | 25.8 | 10.5 | 1.24 | 3.63E-02 |

| | | | | | | | | | |
|---|---|---|---|---|---|---|---|---|---|
| GL16 | UP | 1342 | 5.65 | 10.63 | 7.82 | -43 | 95.8 | 0.66 | 5.83E-02 |
| GL16 | DN | 1209 | 4.93 | 11.03 | 7.8 | -41.2 | 97.5 | 0.43 | 4.72E-02 |
| GL17 | UP | 1229 | 5.9 | 10.71 | 7.47 | -23.9 | 21.2 | 22.97 | 3.27E-03 |
| GL17 | DN | 1007 | 6.9 | 10.43 | 7.6 | -31.4 | 26 | 16.73 | 2.83E-03 |
| GL18 | UP | 1204 | 4.15 | 11.35 | 7.98 | -51.2 | 119 | 2.15 | 6.63E-04 |
| GL18 | DN | 1117 | 5.53 | 11.04 | 8 | -52.7 | 106.3 | 11.17 | 5.35E-04 |
| GL19 | UP | 1177 | 0.13 | 11.23 | 8.69 | -88.5 | 122.4 | 232.67 | 3.2E-04 |
| GL19 | DN | 1094 | 1.8 | 11.25 | 8.29 | -68.3 | 126.5 | 65.53 | 4.2E-04 |
| GL20 | UP | 1475 | 2.63 | 11.8 | 7.83 | -42.9 | 68.4 | 1.46 | 5.76E-04 |
| GL20 | DN | 990 | 5.28 | 11.43 | 7.81 | -41.2 | 62.6 | 2.85 | 9.84E-04 |
| GL21 | UP | 1084 | 3.4 | 11.99 | 7.59 | -29.9 | 33.7 | 2.3 | 2.44E-03 |
| GL21 | DN | 1014 | 2.68 | 12.25 | 7.54 | -26.9 | 25.4 | 4.7 | 6.8E-04 |



m.a.s.l.: meters above sea level, DO: Dissolved Oxygen, NTU: Nephelometric Turbidity Units, n/d: non-determined

**Table S6. The summary of the step-wise model building for the distance-based redundancy analysis.** The "+" sign before each variable indicates its step-wise addition to the model formula.

| | **Cumulative Adjusted $R^2$ (%)** | **Df** | **AIC** | **F** | **p** |
|---|---|---|---|---|---|
| **Conductivity** | 14.14 | 1 | 414.19 | 20.45 | **0.002** |
| **+ Dissolved Oxygen** | 22.25 | 1 | 403.37 | 13.2 | **0.002** |
| **+ Latitude** | 27.99 | 1 | 395.21 | 10.24 | **0.002** |
| **+ Chl-α** | 33.61 | 1 | 386.5 | 10.74 | **0.002** |
| **+ pH** | 37.88 | 1 | 380.5 | 7.86 | **0.002** |
| **+ Water T** | 40.21 | 1 | 375.93 | 6.35 | **0.002** |
| **+ Turbidity** | 42.15 | 1 | 372.94 | 4.76 | **0.002** |
| **All variables** | 43.54 | | | | |

Df: Degrees of freedom, AIC: Akaike Information Criterion
